# Probing *Clostridium difficile* infection in innovative human gut cellular models

**DOI:** 10.1101/269035

**Authors:** Blessing O. Anonye, Jack Hassall, Jamie Patient, Usanee Detamornrat, Afnan M. Aladdad, Stephanie Schüller, Felicity R.A.J. Rose, Meera Unnikrishnan

## Abstract

Interactions of anaerobic gut bacteria, such as *Clostridium difficile*, with the intestinal mucosa have been poorly studied due to challenges in culturing anaerobes with the oxygen-requiring gut epithelium. Although gut colonization by *C. difficile* is a key determinant of disease outcome, precise mechanisms of mucosal attachment and spread remain unclear. Here, using human gut epithelial monolayers co-cultured within dual environment chambers, we demonstrate that *C. difficile* adhesion to gut epithelial cells is accompanied by a gradual increase in bacterial numbers. Prolonged infection causes redistribution of actin and loss of epithelial integrity, accompanied by production of *C. difficile* spores, toxins and bacterial filaments. This 2-D dual chamber system was used to examine *C. difficile* interactions with the commensal *Bacteroides dorei*, and interestingly, *C. difficile* growth is significantly reduced in presence of *B. dorei*. Furthermore, in novel multilayer and 3-D gut models containing a myofibroblast layer, *C. difficile* adheres more efficiently to epithelial cells, as compared to the 2-D model, leading to a quicker destruction of the epithelium. Our study describes new controlled environment human gut models that enable host-anaerobe and pathogen-commensal interaction studies *in vitro*.

## INTRODUCTION

*Clostridium difficile*, an anaerobic spore-forming bacterium, is the main cause of infectious diarrhea in healthcare settings. A major risk factor for *C. difficile* infection (CDI) is antibiotic use, which results in the disruption of the intestinal microbiota, allowing *C. difficile* to colonize and proliferate (Adamu and Lawley, 2013; Fuentes et al., 2014). CDI is one of the most common healthcare-associated infections in the United States, with an estimated 453,000 cases of CDI and 29,000 deaths reported in 2011, and with the highest mortality rates among the elderly (Lessa et al., 2015). Although the number of cases in the UK are declining [13,000 cases in 2016-2017 (England, 2017)], an increasing incidence of CDI has been reported across Europe, Canada and Australia (Gravel et al., 2009; Davies et al., 2014; Collins et al., 2017).

CDI pathogenesis is complex and mediated by a number of bacterial virulence factors. *C. difficile* produces two main toxins, toxin A and toxin B (TcdA and TcdB), and some strains produce a binary toxin (CDT), all of which contribute to bacterial pathogenicity (Kuehne et al., 2014; Carter et al., 2015). Toxins A and B are known to disrupt the intestinal epithelial barrier function through inactivation of the Rho GTPases, which results in reorganization of the actin cytoskeleton and induction of the MAPK pathways, leading to a cytokine response (Bobo et al., 2013; Chen et al., 2015). The key role for these toxins during infection has been demonstrated in animal models using toxin mutants, which are unable to cause disease and death (Lyras et al., 2009; Kuehne et al., 2010; Kuehne et al., 2014; Carter et al., 2015). While toxins are major virulence factors, recent studies have highlighted the importance of several *C. difficile* surface proteins such as surface layer proteins (Calabi et al., 2002; Merrigan et al., 2013), adhesins (Hennequin et al., 2001; Barketi-Klai et al., 2011; Kovacs-Simon et al., 2014), cell wall proteins (Waligora et al., 2001), pili and flagella in CDI (Tasteyre et al., 2001; Batah et al., 2017; McKee et al., 2018). *C. difficile* is also known to produce spores that are resistant to antibiotics and disinfectants (Lawley et al., 2010; Vohra and Poxton, 2011). *C. difficile* spores mediate disease transmission and may serve as a reservoir within the host causing recurrence of infection (Deakin et al., 2012b).

Although animal models of CDI have been used to understand *C. difficile* pathogenesis and investigate the functions of several *C. difficile* factors (Chen et al., 2008; Kuehne et al., 2010; Deakin et al., 2012b; Darkoh et al., 2016), it is challenging to study host-bacterial interactions occurring at the gut mucosal interface *in vivo*. *In vitro* human cell culture models enable molecular and cellular studies on both the host and the pathogen, easier testing of multiple conditions and visualisation of infection dynamics. Infection of human intestinal epithelial cell (IEC) lines has been used to study *C. difficile* pathogenesis but these models have been limited to short time periods as *C. difficile* requires an anaerobic environment for optimal growth (Cerquetti et al., 2002; Janvilisri et al., 2010; Mora-Uribe et al., 2016). A dual environment system such as a vertical diffusion chamber (VDC), which permits growth of the bacteria and IECs in appropriate gaseous environments was used previously by Schüller and Phillips to demonstrate increased adherence of enterohaemorrhagic *Escherichia coli* (EHEC) to polarized IECs in microaerobic compared to aerobic conditions, accompanied by enhanced expression and translocation of EHEC type III secreted effector proteins (Schüller and Phillips, 2010). Similarly, an increase in *Campylobacter jejuni* invasion of IECs was observed under a microaerobic environment in the VDC (Mills et al., 2012). Recently, a VDC was employed to culture *C. difficile* with T84 cells where an anaerobic environment was shown to enhance *C. difficile-*induced cytokine production, compared to aerobic co-culture although this was only studied over a short time course (Jafari et al., 2016).

Models that replicate the physiology and local tissue environment found *in vivo* are ideal for investigating pathogen interactions in IECs. Three-dimensional (3-D) gut models support better epithelial cell growth and differentiation through proteins such as growth factors secreted by underlying cell layers (Morris et al., 2014; Kook et al., 2017). Scaffolds generated from natural or synthetic polymers such as matrigel, collagen and polyethylene terephthalate (PET) have been used to generate the extracellular matrix, which is a major tissue component, and supporting scaffolds can be designed to meet the cell type-specific needs. (Ravi et al., 2015; Kook et al., 2017). More recently, electrospinning has been employed to fabricate fibers from polymers creating a structure similar to the natural fibrous network of the extracellular matrix (ECM) (Morris et al., 2014; Kook et al., 2017). Electrospinning was used to create nanofiber and microfiber scaffolds for optimal 3-D *in vitro* culture of airway epithelial and fibroblast cells (Morris et al., 2014). Fibroblasts play an active role in producing ECM and producing chemokines in response to bacterial infection (Smith et al., 1997).

Three-dimensional gut models have been employed to understand key bacterial and host pathways in a range of pathogens including *C. difficile* (Kasendra et al., 2014; Leslie et al., 2015), *Salmonella* (Barrila et al., 2017), *Cryptosporidium parvum* (DeCicco RePass et al., 2017) and Coxsackievirus B (Drummond et al., 2016). In human intestinal organoids infected with *C. difficile*, bacteria were reported to be viable for up to 12h (Leslie et al., 2015), while a 3-D model using Caco-2 cells grown as cysts in a matrigel monitored the adhesion and translocation of *C. difficile* for 1h (Kasendra et al., 2014). While both studies investigated the role of *C. difficile* toxins in these models, neither was able to follow the infection for a longer period of time due to the lack of an optimal environment for bacterial growth.

In this study, we have determined the infection dynamics of *C. difficile* over an extended time frame using novel human gut epithelium models. We demonstrate an increase in the numbers of adherent *C. difficile* accompanied by production of spores, toxins and bacterial filamentous forms, along with host chemokine production, over 48h in a IEC monolayer VDC model. We demonstrate that this system can be used to study interactions of obligate anaerobes such as the gut commensal, *Bacteroides dorei*, with IECs. Interestingly, we show that *C. difficile* replication is significantly reduced in presence of *B. dorei* on gut epithelial cells. In a complex 3-D model that we developed which contains myofibroblasts on electrospun nanofiber scaffolds, we observed increased bacterial adhesion to the IECs, compared to the 2-D model.

## RESULTS

### Development of a gut cellular model for *C. difficile*

Polarized IECs (epithelial Caco-2 and mucus producing HT29-MTX goblet cells) were cultured on the apical side of a Snapwell insert as shown in Figure 1. Inserts with polarized cell layers were placed in the VDC and experiments were performed by perfusing a mixture of 5% CO_2_ and 95% air in the basolateral compartment for epithelial cell growth and an anaerobic gas mixture (10% CO_2_, 80% N_2_ and 10% H_2_) on the apical side for bacterial growth. After 3h, the cell layers were washed with PBS, followed by the addition of fresh prereduced media (to simulate infection conditions described below). The barrier integrity of IECs was monitored by measuring the transepithelial electrical resistance (TEER). The TEER values demonstrated an intact epithelium in the control IECs incubated within the VDC, although a slight decline was observed over 24h (Figure 2A). However, no significant disruption to the IECs was seen after 24h or 48h by microscopy; actin staining showed that the cytoskeleton of the control cells was intact at 24h and 48h (Figure 2B). Immunofluorescent staining of the Snapwell inserts for MUC2, a major mucus protein produced by goblet cells, showed that a small amount of mucus was produced (Figure S1) in this cell layer which contained 10% goblet cells (9 Caco-2:1 HT29-MTX).

**Figure 1.**
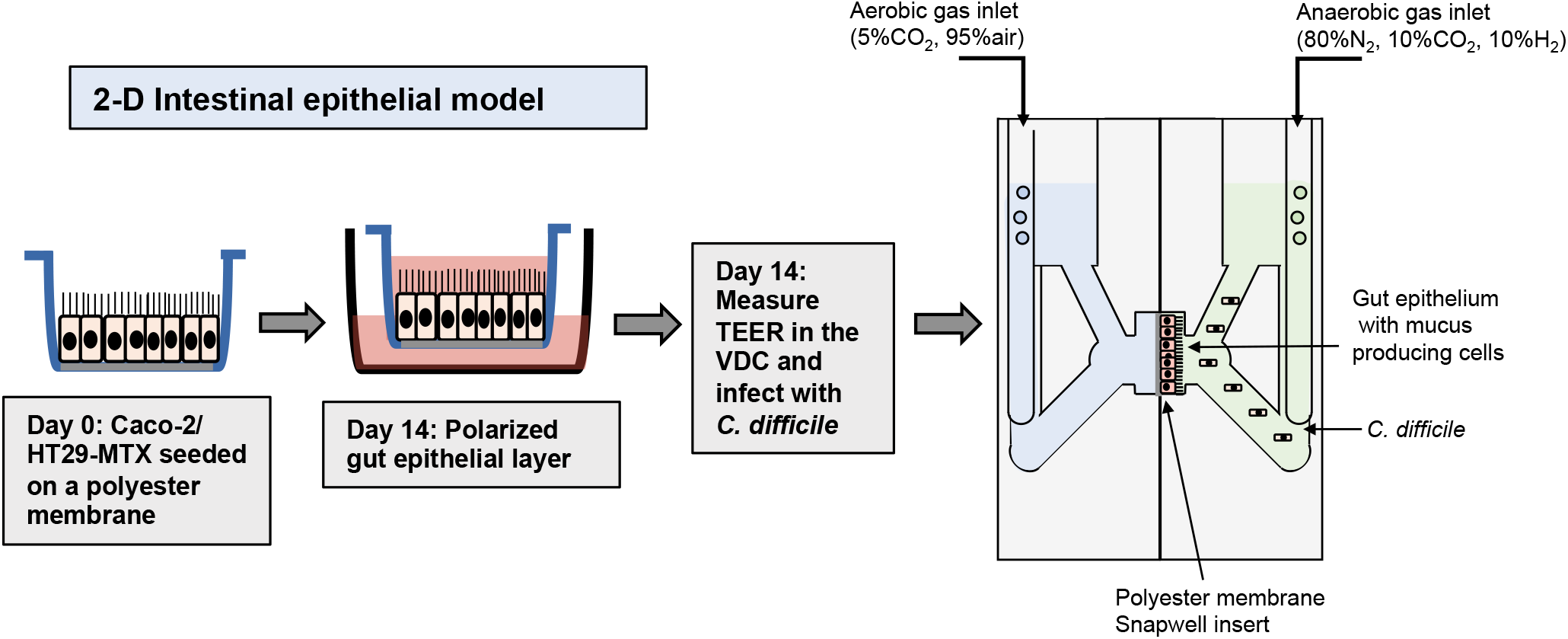
Scheme for generation of epithelial layers for use within vertical diffusion chambers. For the 2-D model, intestinal epithelial cells were grown to form a polarized confluent monolayer in a Snapwell insert for 2 weeks before inserting in the VDCs. *C. difficile* was added to the apical chamber which was perfused with an anaerobic gas mix and incubated for different times while maintaining anaerobic conditions. The basolateral compartment was perfused with a mixture of 5% CO_2_ and 95% air, necessary for growth of IECs.

**Figure 2.**
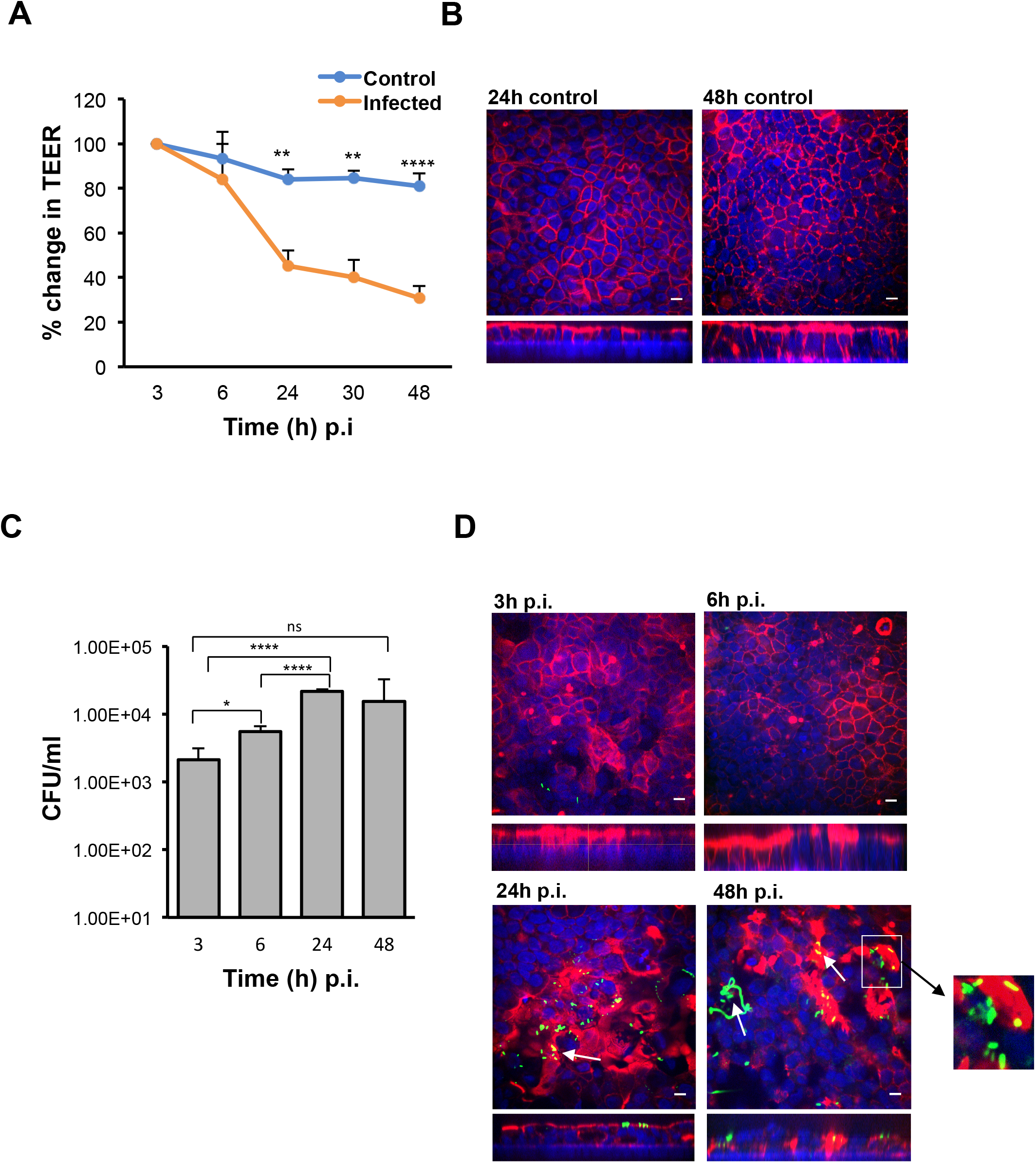
The VDC permits growth of host cells and enables *C. difficile* to adhere to gut epithelial cells. (A) Reduction in TEER measurements at different time after infection indicating increasing permeability compared to uninfected controls, \*\**p* < 0.01, \*\*\*\**p* < 0.0001 as determined by one-way ANOVA. (B) Immunofluorescent microscopy images of uninfected controls at 24h and 48h. Actin is stained with phalloidin (red) and cell nuclei are stained with DAPI (blue), scale bar = 10 µM. (C) Colony counts from infected cell lysates to determine the number of adherent or cell-associated bacteria. A significant increase in adherent *C. difficile* is observed from 3h to 24h. Data shown is the mean of 3 independent experiments in triplicates and error bars indicate SD, \**p* < 0.05 and \*\*\*\**p* < 0.0001 as determined by one-way ANOVA. (D) Immunofluorescent microscopy images of *C. difficile* infected IECs at 3h, 6h, 24h and 48h p.i. showing colocalisation (yellow) of the bacteria stained with anti *C. difficile* antibodies (green) with actin, stained with phalloidin (red). Cell nuclei are stained with DAPI (blue). Reorganization and destruction of actin filaments is observed at 24h and 48h p.i. Inset (white square box) shows micro-communities, and bacterial colocalisation with actin at 24h and 48h (yellow) and *C. difficile* filamentous forms observed at 48h p.i. are indicated by white arrows, scale bar = 10 µM. Inset below shows the orthogonal XZ axis view of the IECs.

Furthermore, to ensure that anaerobic conditions were maintained in the apical chamber, growth of *C. difficile* in the apical compartment of the VDC was compared to growth in the anaerobic cabinet for 3h and 24h. No negative impact on growth was observed at 3h and 24h compared to bacteria grown in the anaerobic cabinet; instead, a slight increase in *C. difficile* growth in the VDC was noted at 24h (Figure S2).

### *C. difficile* colonization leads to disruption of the intestinal epithelium

To determine how *C. difficile* interacts with the human host in the short and long term, Caco-2/HT29-MTX layers were infected with *C. difficile* R20291 at an MOI of 100:1 for different periods of time in the anaerobic chamber of the VDC (Figure 1). In order to study bacteria that adhere to the IECs and their replication, for all experiments, at 3h p.i., the apical supernatant containing the *C. difficile* was removed, the IECs washed in PBS, fresh prereduced media added, followed by incubation for the required time. Uninfected controls shown in Fig 2B were run in parallel. The number of adherent *C. difficile* was determined by counting colony forming units (CFUs) from the cell lysates, after washing off non-adherent bacteria. A significant increase in the number of cell-associated *C. difficile* was observed from 3h to 24h p.i. (Figure 2C). This increase in cell-associated bacteria corresponded to a decrease in TEER measurements indicating disruption of the intestinal epithelial barrier (Figure 2A, Figure S3A). Confocal microscopy showed *C. difficile* present as small micro-communities on the IECs at 24 and 48h (Fig 2D). An additional image of microcommunities formed at 24h is shown in Fig S3B At early time points (3h and 6h p.i.), there was little disruption of the actin filaments but at 24h and 48h p.i., destruction of the cytoskeleton was evident (Figure 2D). Interestingly, at 48h p.i., immunofluorescent staining of bacteria showed the presence of filamenting *C. difficile* (Figure 2D). A Coloc 2 ImageJ colocalisation analysis revealed partial colocalisation of *C. difficile* with actin (Figure 2D) as indicated by the Manders’ M2 value (channel 2 for *C. difficile*) which shows 30% and 20% *C. difficile* colocalised with the actin at 24h and 48h p.i respectively (Table 1) Positive Li’s ICQ values and Costes signifance test values (Table 1) further confirmed colocalisation at both time points.

**Table 1.**
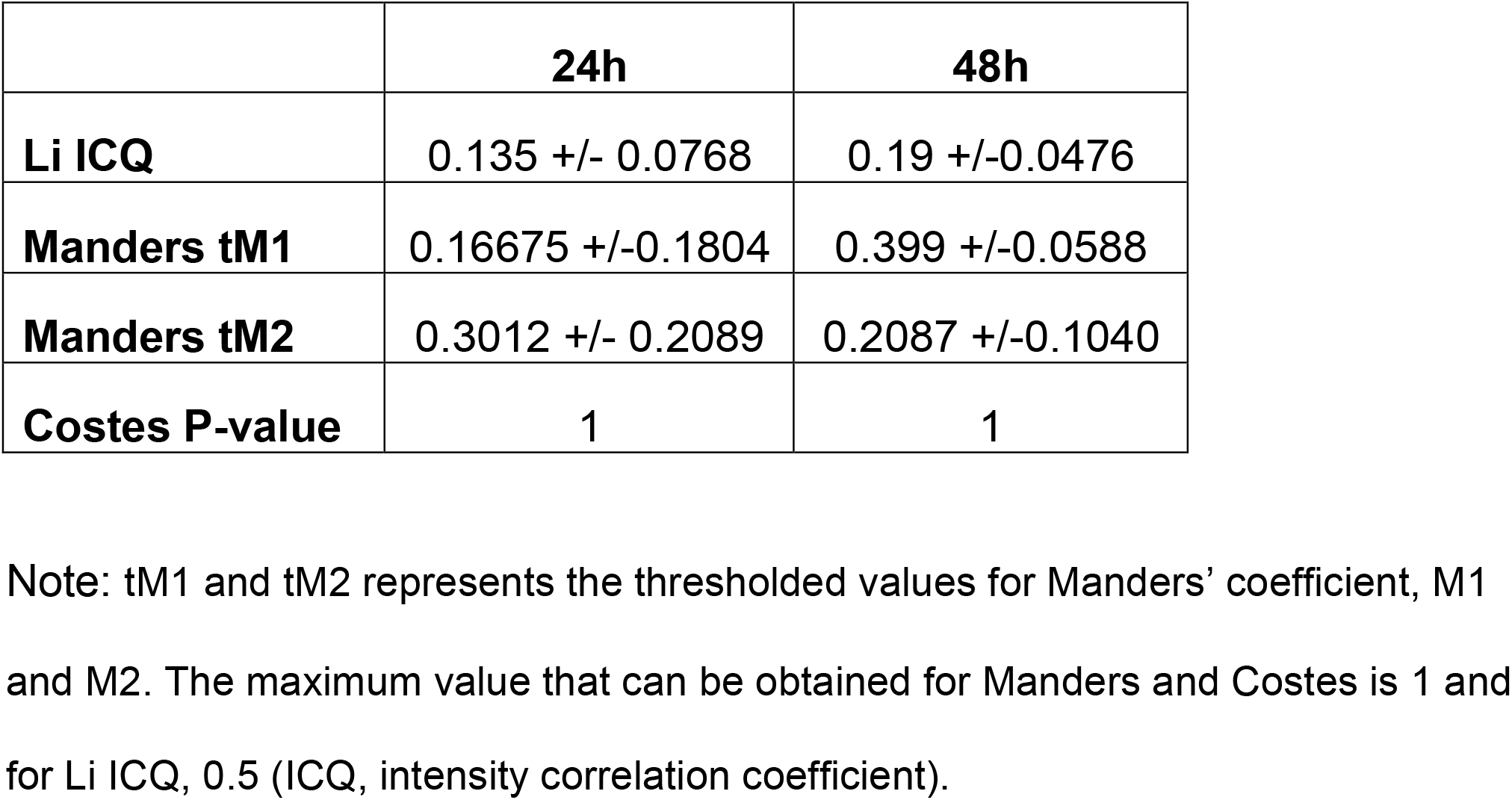
Colocalisation analysis using ImageJ.Coloc 2.

|  | **24h** | **48h** |
| --- | --- | --- |
| **Li ICQ** | 0.135 +/- 0.0768 | 0.19 +/-0.0476 |
| **Manders tM1** | 0.16675 +/-0.1804 | 0.399 +/-0.0588 |
| **Manders tM2** | 0.3012 +/- 0.2089 | 0.2087 +/-0.1040 |
| **Costes P-value** | 1 | 1 |

Note: tM1 and tM2 represents the thresholded values for Manders' coefficient, M1 and M2. The maximum value that can be obtained for Manders and Costes is 1 and for Li ICQ, 0.5 (ICQ, intensity correlation coefficient).

### Prolonged infection is associated with spore and toxin production, and host responses

In order to fully understand bacterial factors necessary for CDI persistence, we studied the production of spores, toxins and a host chemokine in the 2-D monolayer epithelial model. We measured spores and total bacteria from the cell-associated fraction at different times after infection. Although there were ∼0.1% spores in the inoculum, there was a mild but significant increase in spore numbers from 3h to 48h p.i. (*p* < 0.05, Figure 3A). After washing off the non-adherent bacteria at 3h, we also tracked the bacterial spore numbers from 3h-48h in the supernatants (Figure S4). By 48h p.i., there were equal numbers of spores and total cells in the supernatants (Figure S4).

**Figure 3.**
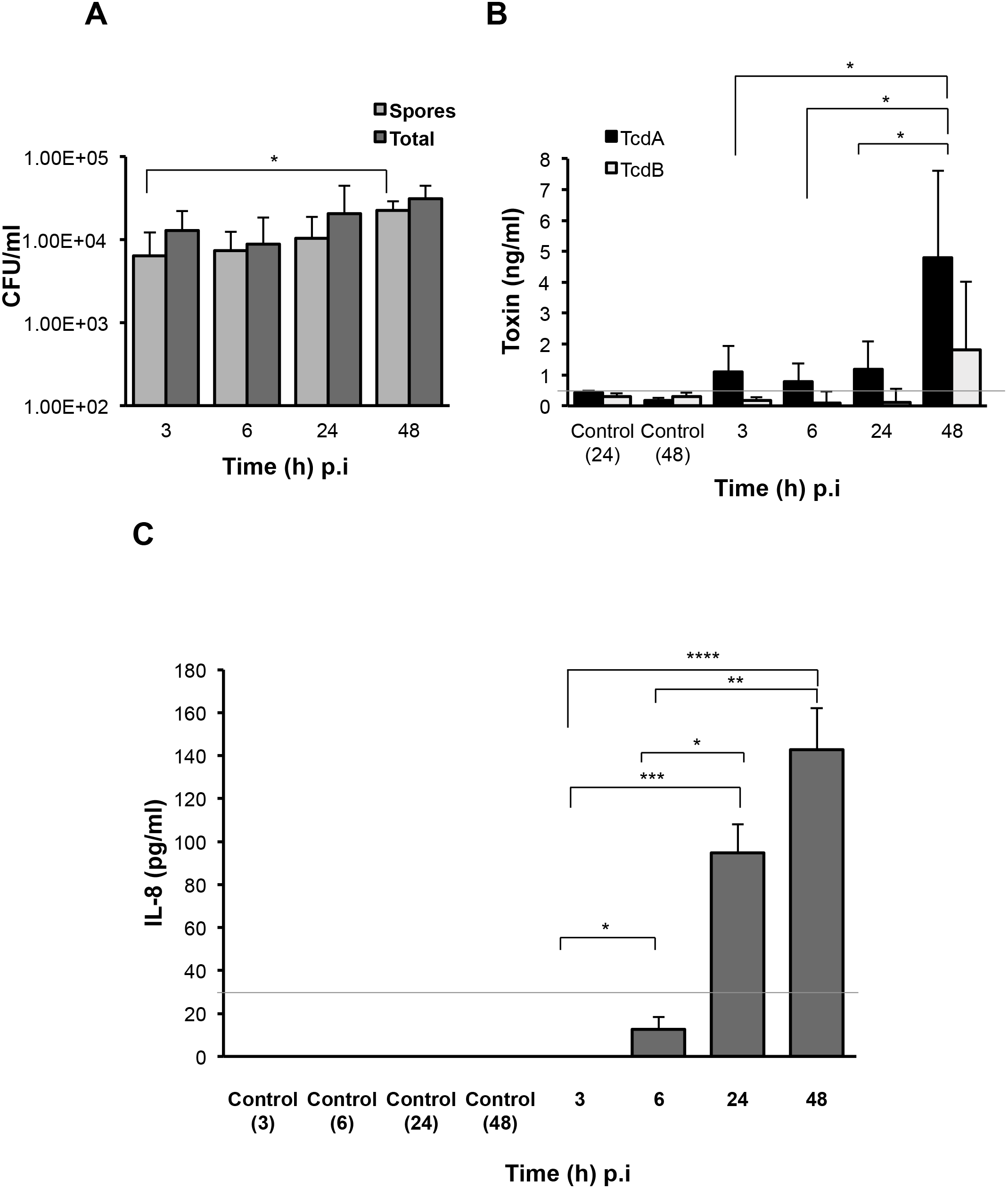
Production of *C. difficile* spores, toxins and host responses to infection. (A) Colony counts of spores recovered after heat treatment, and total cells in the cell-associated *C. difficile* fraction (infected cell lysates) in the 2-D epithelial model. Data shown is the mean of 3 independent experiments and error bars indicate SD, \**p* < 0.05 as determined by two-way ANOVA (B) ELISA for *C. difficile* toxins A and B shows increased toxin production after extended infection. Toxins were measured from medium obtained from the apical compartment containing uninfected cell layers incubated for 24h or 48h (Control) or cells infected with *C. difficile* for 3h, 6h, 24h or 48h. Data shown is the mean of 3 independent experiments and error bars indicate SD, \**p* < 0.05 as determined by two-way ANOVA. Grey line represents the sensitivity of the test at 0.5 ng/ml. (C). ELISA for human IL-8 indicates increased IL-8 production at 24h and 48h p.i. IL-8 was measured in medium obtained from the basolateral compartment containing uninfected cell layers incubated for 3h, 6h, 24h or 48h (Control) or cells infected with *C. difficile* for 3h-48h. Grey line represents the limit of detection at 32 pg/ml. Data shown is the mean of 3 independent experiments and error bars indicate SD, \**p* < 0.05, \*\**p* < 0.01 and \*\*\**p* < 0.001, *p* < 0.0001 as determined by one-way ANOVA.

Toxins A and B (TcdA and TcdB) were monitored over time during infection using ELISA. Toxin A increased significantly from 3h to 48h p.i. (*p* < 0.05) while low levels of toxin B were detected at all times (Figure 3B). Analysis of the basolateral compartment supernatant for the host chemokine IL-8, which has been previously implicated in *C. difficile* infection (Rao et al., 2014), revealed that IL-8 levels were low at 6h p.i. but increased at 24h and 48h p.i. in this infection model compared to uninfected controls (Figure 3C).

### Co-culturing *C. difficile* with *Bacteroides dorei* results in reduced *C. difficile* growth within an epithelial gut model

To determine if our VDC-based system could be extended for use with other anaerobic bacteria, we studied a strict commensal anaerobe, *B. dorei* within the epithelial monolayer model using similar conditions as for *C. difficile*. Adherence to the IECs was determined at 3h and 24h by CFU counts, as described for *C. difficile*. *B. dorei* adhered to the epithelial cells at 3h and multiplied over 24h, as observed for *C. difficile* (Figure 4 A, B). A mixed culture of *B. dorei* and *C. difficile* (1:1), prepared as described in Experimental Procedures was then cultured with the monolayer in the VDC. Cell-associated bacteria were quantitated by plating on a medium used to selectively isolate *C. difficile* colonies, which also allowed the growth of *B. dorei*. *Bacteroides* colonies were distinguished by colony size, color and morphology (small colonies). A significant decrease in the number of *C. difficile* was observed when co-cultured with *B. dorei* in the presence of IECs when compared to mono-cultures of *C. difficile* at 24h p.i., but not at 3h p.i. (Figure 4A, B, *p* < 0.001). We also observed higher colony counts when *B. dorei* was grown in co-culture with *C. difficile* at 3h and 24h, compared to mono-cultures of *B. dorei* (Figure 4A, B).

**Figure 4.**
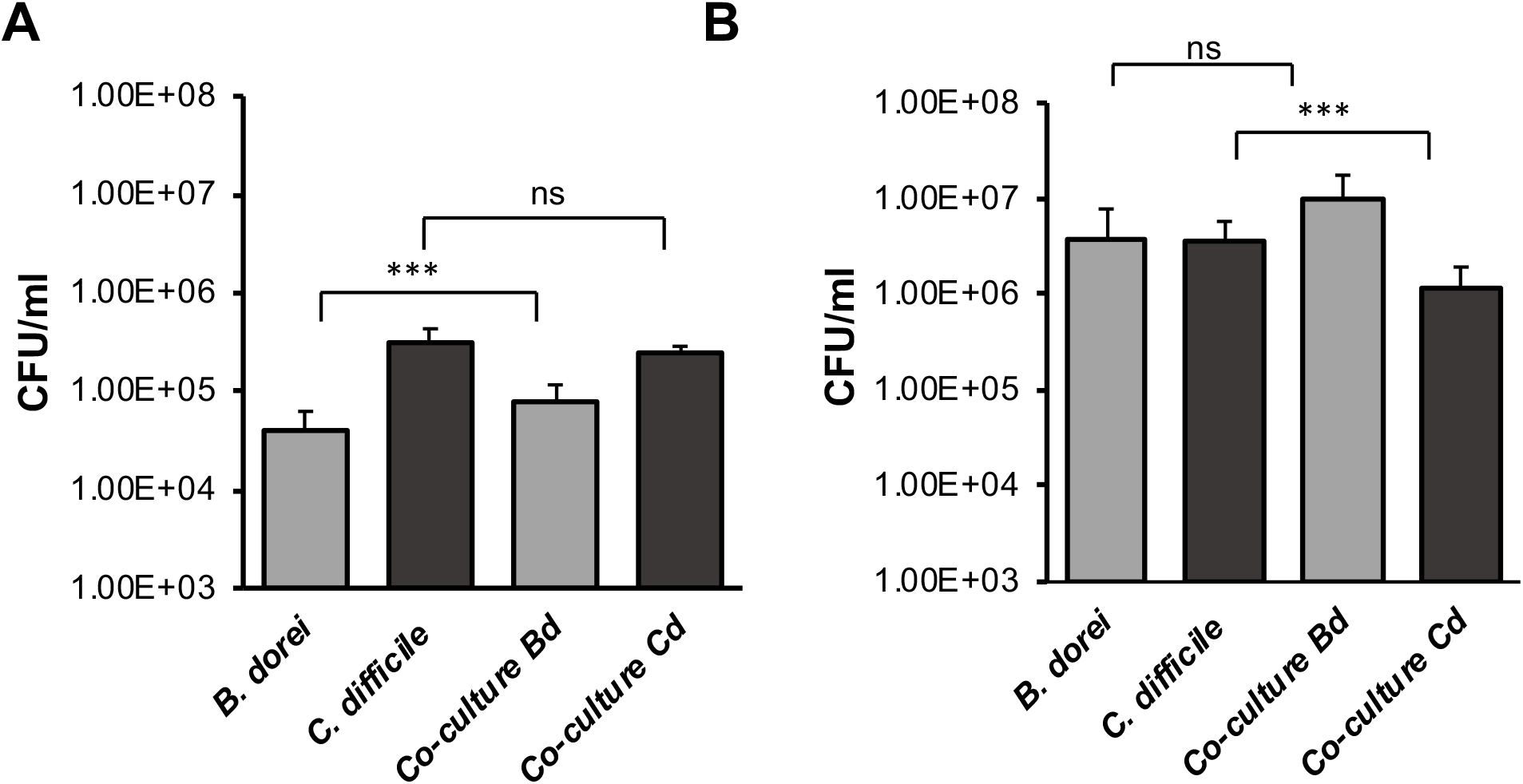
Studying *C. difficile*-commensal interactions within epithelial models. (A) Mono or mixed cultures of *B. dorei* and *C. difficile* were incubated with the epithelial monolayers in VDCs for 3h. Co–cultures of *B. dorei* with *C. difficile* in the VDC showed no significant reduction of *C. difficile* growth at 3h p.i. as compared to *C. difficile* mono-culture. A relative increase in *B. dorei* adhering to the IECs was observed in co-cultures. ‘Co-culture Bd and Cd’ represents the growth of *B. dorei* and *C. difficile* respectively in the mixed culture (Bd, *B. dorei* and Cd, *C. difficile*) on the IECs. Data shown is the mean of 3 independent experiments and error bars indicate SD, \*\*\**p* < 0.001, ns, not significant as determined by one-way ANOVA. (B) At 24h, a significant reduction of *C. difficile* CFU is observed in co-cultures compared with *C. difficile* mono-culture. Data shown is the mean of 3 independent experiments and error bars indicate SD, \*\*\**p* < 0.001, ns, not significant as determined by one-way ANOVA.

### Increased cell-associated bacteria in multilayer and 3-D models

Typically, a myofibroblast layer underlies the basement membrane in the human gut. To develop this system further by increasing its cell complexity, we incorporated myofibroblasts into our model. Human CCD-18co myofibroblasts were first grown on the basolateral side of the polyester Snapwell insert before seeding the IECs on the apical side (multilayer model, Figure 5A). Furthermore, to recreate the highly porous architecture of the ECM in the basement membrane, epithelial and fibroblasts cells were grown apically and basolaterally respectively on inserts containing electrospun nanofiber scaffolds generated from polyethylene terephthalate to develop a 3-D gut epithelium as reported previously (Morris et al., 2014). Scanning electron microscopy revealed that these scaffolds exhibited a uniform nanofibrous matrix (Figure 5B) that supported the attachment and proliferation of Caco-2 cells to form a confluent monolayer (Figure 5C) and enabled the proliferation of the CCD-18co cells (Figure 5D). The average fiber diameter was 457 ± 170 nm (Figure S5).

**Figure 5.**
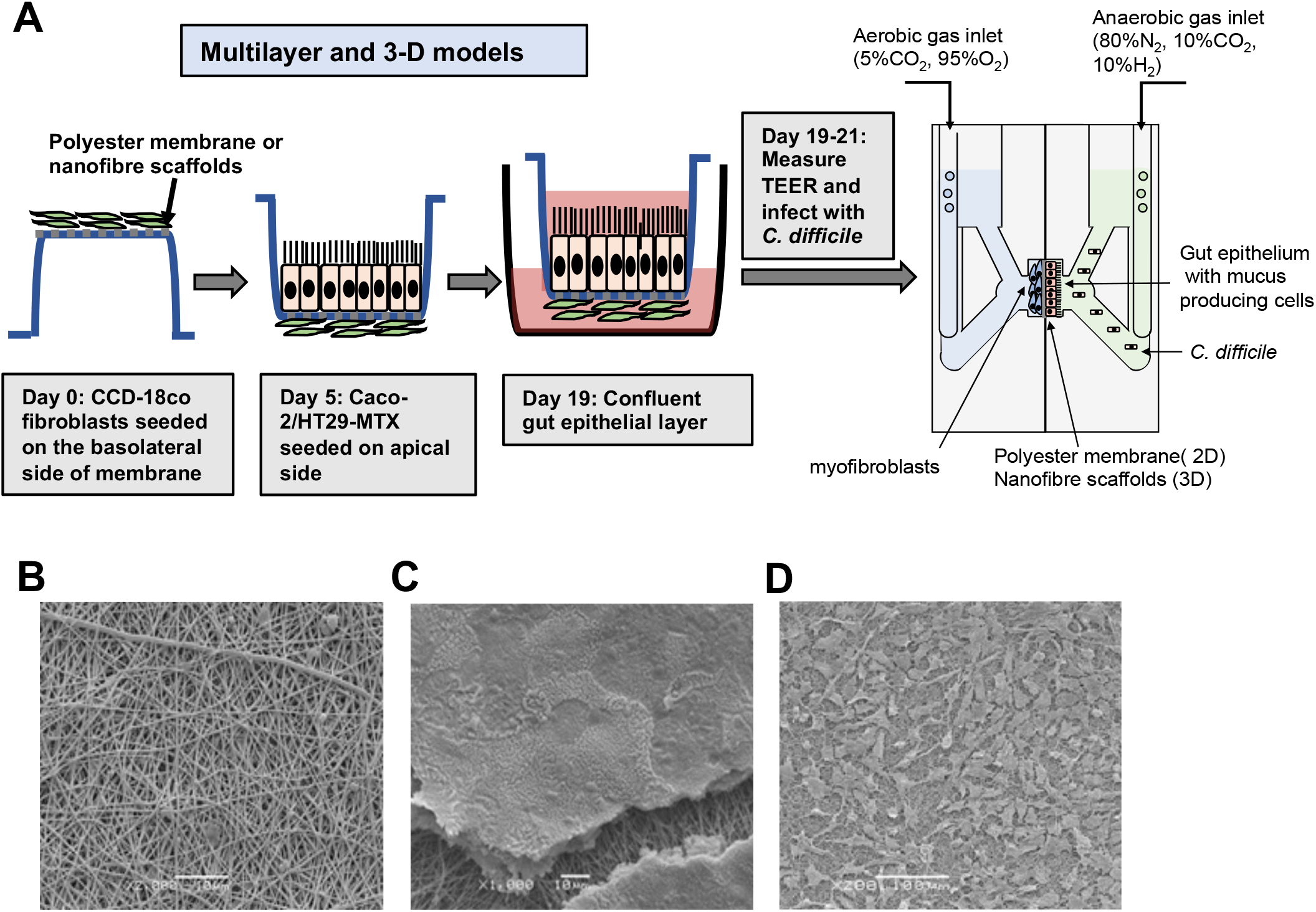
Scheme for generation of multilayer and 3-D models containing fibroblast cells and cell growth on the electrospun nanofibrous matrix. (A) Fibroblast cells were first seeded on the basolateral side of the Snapwell or nanofiber scaffold for the multilayer and 3-D models for 5 days. Thereafter, IECs were cultured on the apical side for 2 weeks before placing the Snapwell insert between two halves of the chamber. *C. difficile* was infected apically while maintaining anaerobic conditions and the basolateral compartment was perfused with 5% CO_2_ and 95% air. (B) Polyethylene terephthalate (PET; 10% w/v) was electrospun and exhibited a uniform nanofibrous matrix as determined by scanning electron microscopy. (C) Caco-2 epithelial cells were able to form a confluent monolayer on the nanofibrous scaffolds (the scaffold can be seen underneath the cells following processing for SEM). (D) The CCD-18co cells proliferated across the nanofibrous scaffold forming cell processes to adhere to the nanofibers.

Interestingly, in the multilayer model, we find that after infection with the same *C. difficile* strain, same MOI’s (100:1) and conditions as the monolayer model, for 3h and 24h (Figure 2), higher numbers of bacteria adhered to the IECs at both time points (Figure 6A) compared to the monolayer model without fibroblasts (Figure 2). Anti-fibronectin staining of the basolateral side of the membrane containing the fibroblast cells indicated likely degradation of fibronectin, as indicated by the destabilsed fibronectin network, and damage to the fibroblast layer at 24h p.i. (Figure 6B). As before, immunofluorescent staining showed the localisation of *C. difficile* on the epithelial cells at 3h and 24h p.i. (Figure 6C). Infections were not followed for longer times (48h) as the cell layer was badly damaged by 24h p.i.

**Figure 6.**
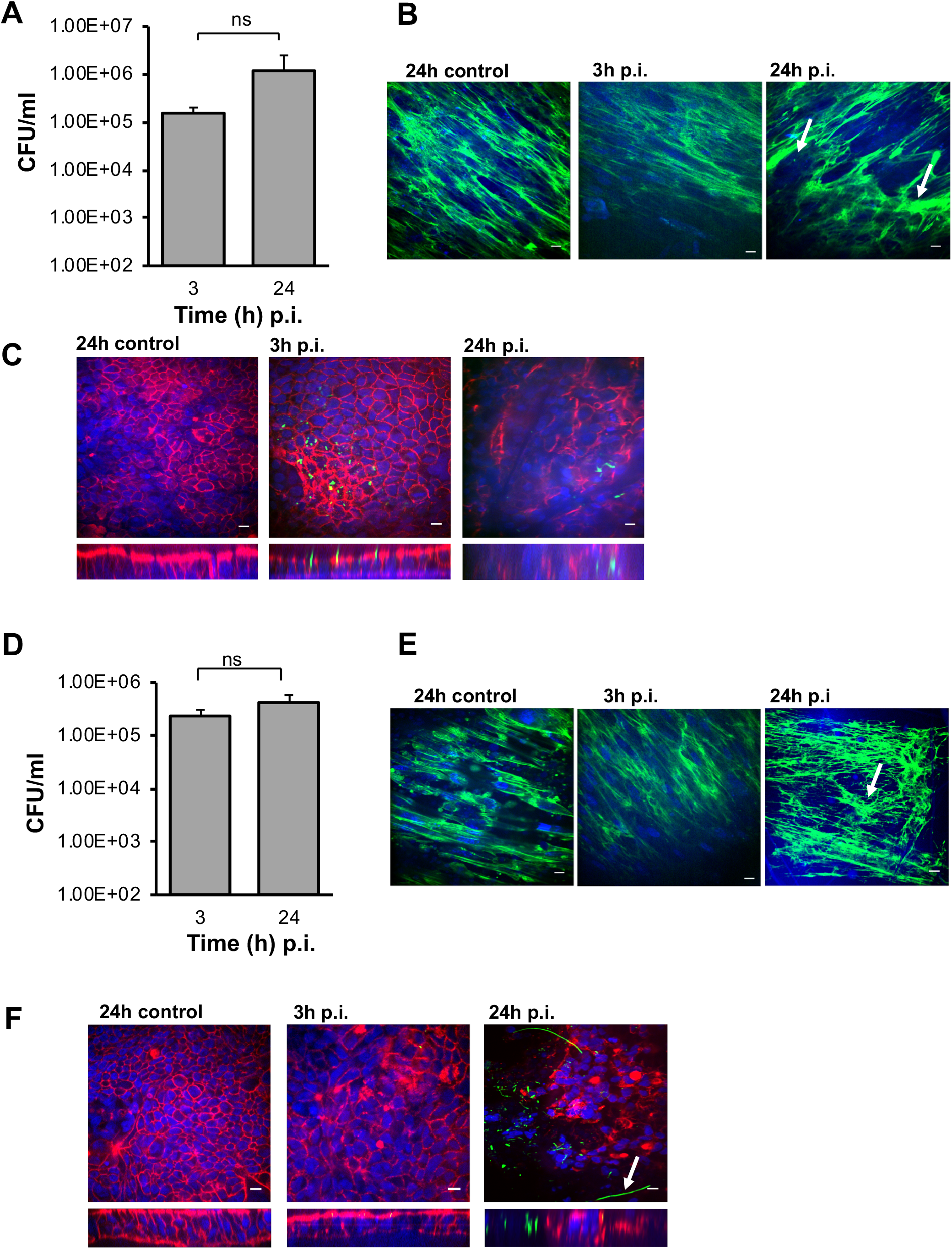
Infection of *C. difficile* in multilayer and 3-D model leads to increased adhesion. Increased adherence of *C. difficile* in the multilayer and 3-D models, where fibroblasts are grown as a basolateral layer on polyester and nanofibers respectively. (A) Serial dilutions of the infected epithelial cell lysates were plated to determine the number of cell-associated bacteria in the multilayer model. Data shown is the mean of 3 independent experiments and error bars indicate SD, ns, not significant as determined by student t test (B) Anti-fibronectin (green) staining of fibroblast cells in the basolateral layer of the control at 24h p.i and infected cells at 3h and 24h p.i. showing redistribution of fibronectin, damaged fibroblast layer in the multilayer model, as indicated by a white arrow. (C) Representative images of *C. difficile* on the apical layer at 3h and 24h p.i. Inset below shows the orthogonal XZ axis view of the IECs. *C. difficile* is stained green, actin, red and cell nuclei, blue, Scale bar = 10 µM. (D) *C. difficile* colony counts from the 3-D model infected cell lysates. Data shown is the mean of 3 independent experiments and error bars indicate SD, ns, not significant as determined by student t test (E) Anti-fibronectin (green) staining of fibroblast cells in the basolateral layer at 3h and 24h p.i. (F) Immunofluorescent microscopy images of *C. difficile* infected IECs at 3h and 24h p.i. in the 3-D model. *C. difficile* filamentous forms are observed at 24h p.i. (indicated by arrow). Inset below shows the orthogonal XZ axis view of the IECs. *C. difficile* is stained green, actin, red and cell nuclei, blue, scale bar = 10 µM.

Similar to the multilayer model, we observed higher adhesion of *C. difficile* to the IECs at 3h and 24h p.i. in the 3-D gut epithelium (Figure 6D) compared to the 2-D model data in Figure 2. Similarly, anti-fibronectin staining showed a destabilisation of the discrete fibronectin network and likely cellular damage of the fibroblast layer in the 3-D model, as indicated by the lack of discrete nuclear staining (Figure 6E). Confocal microscopy revealed the presence of numerous *C. difficile* on the IECs at 24h p.i. (Figure 6F). Interestingly, at 24h p.i., immunofluorescent staining showed the presence of filamenting *C. difficile* (Figure 6F), much earlier than seen in the monolayer infection model. Bacterial staining did not always correlate to the CFU counts (Figure 6C and 6F), which we attribute to the loss of attached bacteria during the staining procedure.

### Spore, toxin and host chemokine production in response to *C. difficile* in 3-D and multilayer models

In both the 3-D and multilayer models of infection, spores were found to adhere to the epithelium at numbers comparable to the 2-D monolayer model (Figure 7A, Figure 3A, Figure S6A), although there was no increase in spore numbers over time. A higher increase in total cell numbers was observed over time in the 3-D and multilayer model (Figure 7, Figure S6A), unlike the monolayer model (Figure 3A). While there was higher variability, levels of toxin A in the apical compartment supernatants from both models were comparable to that produced in the 2-D models (Figure 7B, Figure S6B). Levels of toxin B detected in these models were also low (Figure 7B, Figure S6B). Levels of IL-8, increased over time (3-24h p.i.) in the 3-D (Figure 7C), and multilayer models (Figure S6C), although these were not significantly different to the uninfected controls incubated for 3h or 24h within the VDCs.

**Figure 7.**
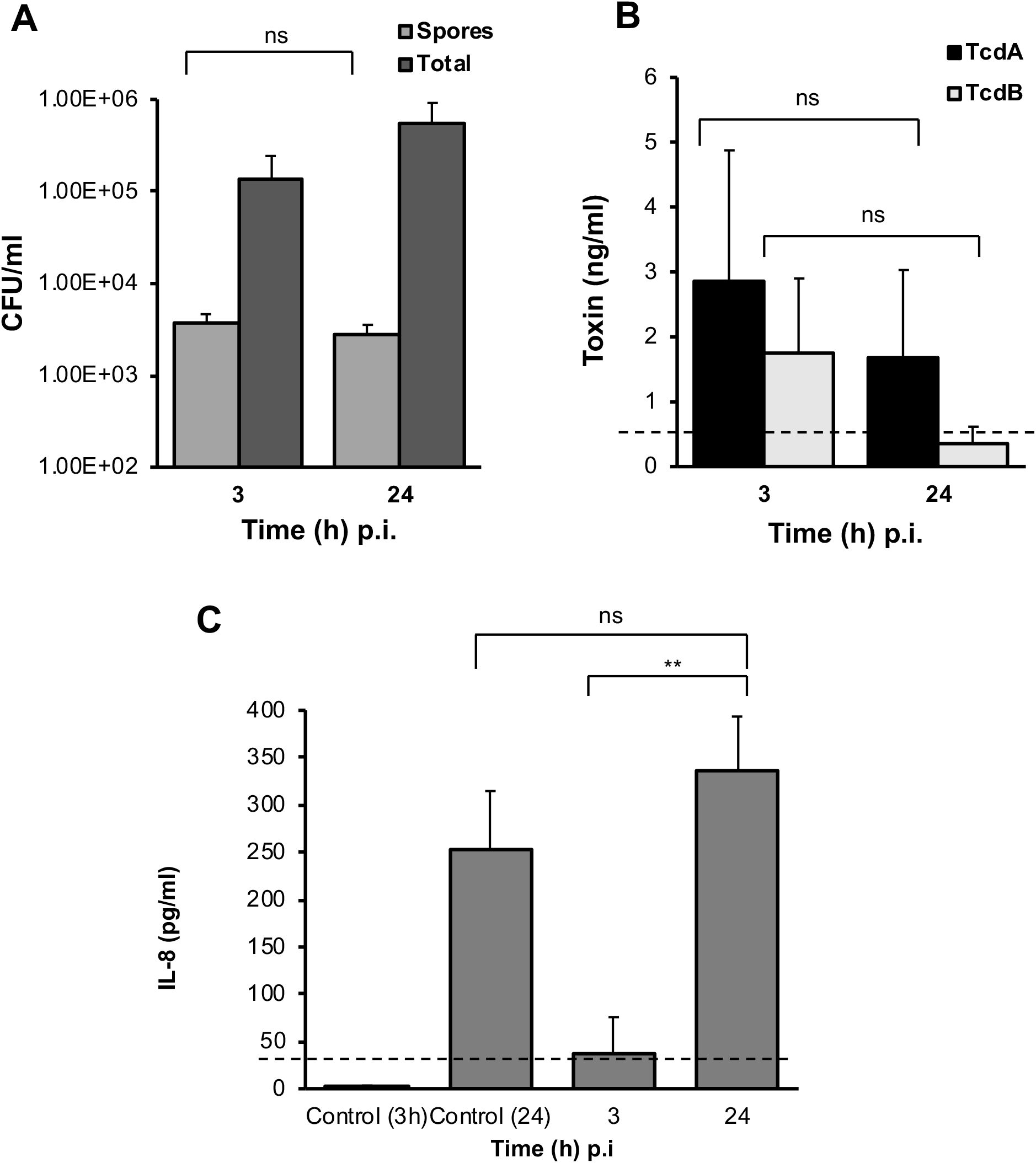
*C. difficile* spores and toxin production, and host response to infection in the 3-D model. (A) Colony counts of spores and total cells in the host cell-associated *C. difficile* fraction (infected cell lysates). Data shown is the mean of 3 independent experiments and error bars indicate SD, ns, not significant as determined by two-way ANOVA. (B) Toxin A and B levels from apical compartment supernatants as determined by ELISA in the 3-D model. Data shown is the mean of 3 independent experiments and error bars indicate SD, ns, not significant as determined by two-way ANOVA. Grey line represents the sensitivity of the test at 0.5 ng/ml (C) Human IL-8 levels in supernatants from the basolateral compartments with uninfected cells incubated for 24h or with cells infected with *C. difficile* for 3h and 24h, as determined by ELISA. Grey line represents the limit of detection at 32 pg/ml. Data shown is the mean of 3 independent experiments and error bars indicate SD, \*\**p* < 0.01. as determined by the one-way ANOVA.

## DISCUSSION

Attachment to the intestinal mucosa, subsequent multiplication and penetration of the gut epithelium are generally essential for the establishment of a successful invasive infection (Kalita et al., 2014; Ribet and Cossart, 2015). We report here the development of 2-D and 3-D *in vitro* human gut models for studying interactions of obligate anaerobes with the host gut epithelium.

We have tracked *C. difficile* infection events over a prolonged time frame in an *in vitro* dual environment human gut model. Along with demonstrating *C. difficile* adhesion, micro-communities, toxin and spore production, we report formation of *C. difficile* filaments, a potential adaptation mechanism during infection.

Previous *in vitro* infection models have shown the attachment of *C. difficile* on IECs either using conditions that are not appropriate for the bacterium (such as growing in aerobic conditions) or that are limited to short term infection due to the anaerobic growth requirements of *C. difficile* (Cerquetti et al., 2002; Paredes-Sabja and Sarker, 2012; Jafari et al., 2016). However, in order to study host-pathogen interactions at a molecular and cellular level, precise environmental control and longer scales of infection are essential. The monolayer (2-D) gut epithelial model described here is ideal for investigating molecular mechanisms underlying *C. difficile*-epithelial interactions as it offers the ability to make cellular, immunological and molecular measurements along with easy substitution of knockout cell lines and introduction of additional cellular components like immune cells.

In the monolayer model, an increase in spores attached to the host cells was observed at 48h p.i. indicating that spore formation occurs during *C. difficile* infection of the gut epithelium in our monolayer model, as reported previously using mouse models (Deakin et al., 2012a). While we cannot be sure if spores are formed and then adhere or if the vegetative cells adhere to the gut cells and sporulate, our findings clearly support previous studies indicating spore adhesion to gut cells during infection. *C. difficile* spores were reported to adhere to undifferentiated Caco-2 cells after 1h of infection as determined by viable spore counts and fluorescence microscopy (Paredes-Sabja and Sarker, 2012), and recently, a spore surface protein CotE was shown to be essential for spore binding to mucus producing epithelial cell layers (Hong et al., 2017).

Toxin production by *C. difficile* is known to play a role in pathogenesis by disrupting the barrier integrity of the intestinal epithelium leading to increased permeability and re-organization of actin (Aktories et al., 2017). Surprisingly, although there is a decrease in TEER and actin reorganization, we detect very low levels of toxins at early time points of infection (6-24h). It is possible that low toxin levels, particularly of toxin B, are sufficient to cause loss of the membrane integrity, as they are augmented by other secreted enzymes produced by *C. difficile*. Although a partial bacterial colocalisation with actin was observed later during infection (24 and 48h) we do not at present understand if *C. difficile* mediates any direct interactions with the acin cytoskeleton. The formation of *C. difficile* filaments seen at 24 and 48 p.i. may be associated with host cell contact and infection, as our studies indicate that incubation in similar conditions (conditioned medium from infected cells or growth in DMEM-10 for 48h) in the absence of cells does not induce this phenotype (Figure S8A and S8B). *C. difficile* filaments may be important during infection, as seen for other gut bacteria (Justice et al., 2008), although this needs to be investigated further.

More than 90% of the bacteria inhabiting the gut are obligate anaerobes but little is known about the adhesion properties of individual bacteria to the intestinal epithelium. Previous research on *Bacteroides fragilis*, a dominant bacterium isolated from intestinal tract infections showed that bile salts enhance the adhesion to the intestinal epithelium and biofilm formation (Pumbwe et al., 2007). However, for other dominant members of the Bacteroidetes phylum, there is insufficient data as to how they interact with the gut epithelium. Our studies with *B. dorei*, a common gut anaerobe in healthy individuals, demonstrates bacterial adhesion and multiplication on IECs, and validates the utility of this *in vitro* model in studying host cell interactions of other anaerobic gut bacteria.

While the gut microbiota is known to protect against *C. difficile* infection (Fuentes et al., 2014), the direct interactions between commensals and gut pathogens have been poorly understood. Although many genome sequencing studies have associated members of the Bacteroidetes with healthy status, the roles of individual *Bacteroides* species in the gut remains unclear (Qin et al., 2010; Huttenhower et al., 2012; Lloyd-Price et al., 2016). Recently, it was shown that *Bacteroides ovatus* inhibited *C. difficile* growth in the presence of bile acids (Yoon et al., 2017). In this study, we demonstrate suppression of gut cell-associated *C. difficile* growth by *B. dorei*, in the absence of bile acids and in a physiologically relevant setting. Indeed, the reduction in *C. difficile* may be because *B. dorei* competes better for nutrients, rather than a direct inhibition. In addition to host-pathogen interactions, the models developed in this work allow three-way interaction studies between commensals, pathogens and the host, i.e. the role of commensals in modulating effects of pathogenic bacteria on the human host. Additionally, as these results may also suggest the potential of non-spore forming anaerobic bacteria in suppressing *C. difficile* growth, such *in vitro* tools may have value in screening and developing novel microbial therapies for treatment of *C. difficile* and other anaerobic infections.

Three-dimensional models are known to promote more *in vivo*-like cell proliferation, growth differentiation and cell-to cell contact (Kook et al., 2017). A recent study reported a 3-D intestinal tissue model for *C. difficile*, where the authors demonstrated that *C. difficile* toxin activity was higher in comparison to a 2-D transwell model (Shaban et al., 2018). They report that spores can germinate into vegetative cells within this model, and that vegetative cells can survive upto 48h, although the ability of *C. difficile* spores to germinate was similar in both their models. While both their 3-D model and our models have epithelial and fibroblast cells, albeit with different scaffolds, a key difference is the ability to control the apical and basolateral side environments using the VDC, which may be important when performing sensitive molecular studies. Similar to Shaban et al, we show bacterial replication upto 48h, although we also track bacteria that adhere to the epithelial layer. Additionally, we have a nanofiber scaffold incorporated between the epithelial and fibroflast layers, which creates a porous architecture similar to the basement membrane underlying the gut epithelium, along with myofibroblast cells in the basal layer.

Previously, Morris *et al* showed that nanofibers were beneficial for epithelial cell differentiation but not penetrable to fibroblasts (Morris et al., 2014). In addition to producing an ECM which provides structural and biochemical support to cells, fibroblasts are also known to produce chemokines when activated by bacteria (Smith et al., 1997). The increased *C. difficile* adhesion observed in the multilayer and the 3-D models, which presumably leads to quicker progressing infection, could be the result of indirect modulation of the epithelial barrier by the myofibroblast cells, though at present we do not understand the underlying mechanisms. The increased amount of the chemokine IL-8 observed in the uninfected 3-D and multilayer models could be attributed to the myofibroblasts, although this chemokine response is not specific to *C. difficile* infection. It is interesting to note that inspite of having higher numbers of bacteria attaching in these models, there was no significant increase in the toxin levels. While we do not observe an increase in spore numbers or distinct toxin production profiles in the 3-D and multilayer models, this may be due to the quick progression of infection due to higher bacterial adhesion. We are currently optimising these models using lower MOI’s to enable longer infection experiments.

Both the multilayer and 3-D gut models have huge potential in studying *C. difficile* pathogenesis, particularly for investigating host invasion and interactions with basolateral surface of the host epithelium and are being utilized for studying functions of secreted *C. difficile* factors in the laboratory. Overall, we have developed highly useful tools for studying *C. difficile* host-interactions which we expect to have broad applications in studying anaerobic gut commensals and their interactions with pathogens and the host.

## 4 EXPERIMENTAL PROCEDURES

### 4.1 Bacterial strains and growth conditions

*C. difficile* R20291 strain B1/NAP1/027 R20291 (isolated from the Stoke Mandeville outbreak in 2004 and 2005), was cultured on brain heart infusion (BHI) agar (Sigma-Aldrich, UK) supplemented with 1 g/L cysteine (Sigma-Aldrich, UK), 5 g/L yeast extract under anaerobic conditions (80% N_2_, 10% CO_2_, 10% H_2_) in a Don Whitley workstation (Yorkshire, UK).

### 4.2 Cell culture, media and conditions

Intestinal epithelial cell line, Caco-2 (P6-P21) from American Type Culture Collection, mucus producing cell line, HT29-MTX (P45-P56), gift from Nathalie Juge, Quadram Institute, Norwich, and fibroblast cells, CCD-18co (P10-P17) were used. Caco-2 cells were grown in Dulbecco’s Modified Eagle Medium (DMEM) supplemented with 10% fetal bovine serum (FBS, Labtech, UK), and 1% penicillin-streptomycin (10,000 units/mL penicillin, 10 mg/mL streptomycin, Sigma-Aldrich, UK). HT29-MTX was grown in DMEM and CCD-18co in Eagle’s Minimum Essential Medium media. Both media were supplemented with 10% FBS, 1% penicillin-streptomycin, 2 mM glutamine and 1% non-essential amino acids (Sigma-Aldrich, UK). All cell lines were maintained in 5% CO_2_ in a humidified incubator at 37°C and free from mycoplasma contamination as determined by the EZ-PCR Mycoplasma kit (Biological Industries, USA).

For the epithelial 2-D models, Caco-2 and HT29-MTX were mixed in a 9:1 ratio and 2 × 10^5^ cells/ml were seeded on a 12 mm Snapwell inserts (tissue culture treated polyester membrane, Corning, New York, USA) supported by a detachable ring for 2 weeks to form a polarized monolayer. For the multilayer and 3-D models, CCD-18co (1 X10^4^ cells/ml) were first seeded on the basolateral layer of the polyester Snapwell insert or electrospun nanofiber scaffold for 5 days after which Caco-2 and HT29-MTX were seeded on the apical side as in the 2-D models for 14 days. Prior to the infection experiments, the cell culture medium in the Snapwell inserts was replaced with antibiotic-free medium.

### 4.3 Vertical diffusion chamber set up and measurement of transepithelial electrical resistance

The Snapwell inserts containing the polarized cell layer were placed between the two half chambers of the vertical diffusion chamber (Harvard Apparatus, Cambridge, UK) and sealed with the provided clamps. 2.7 ml DMEM with 10% FBS (DMEM-10) was placed on both sides of the chamber. Transepithelial electrical resistance measurements was performed using Harvard/Navicyte electrodes on the EC-800 Epithelial Voltage Clamp (Harvard Apparatus, Cambridge, UK) over 24h.

### 4.4 Infection of intestinal epithelial cells (IECs) in 2-D, 3-D and multilayer models

A single bacterial colony was inoculated in pre-reduced BHI broth (Oxoid, UK) supplemented with 1 g/L cysteine (Sigma-Aldrich, UK) and 5 g/L yeast extract and incubated at 37°C overnight. The culture was centrifuged at 10,000 g for 5 mins (Eppendorf 5810R) and bacterial pellet was resuspended in DMEM-10 and incubated at 37°C for at least an hour. Bacterial counts determined from this culture were confirmed to be 2 × 10^7^-3 × 10^7^ CFU/ml for every experiment. This culture was used to infect the IECs at a predetermined MOI of 100:1 in the apical side of the VDC containing the IECs. The apical chamber was diffused with anaerobic gas mixture (10% CO_2_, 10% H_2_, 80% N_2_, BOC, UK) and the basolateral compartment with 5% CO_2_ and 95% air (BOC, UK). At 3h p.i., the apical media containing the *C. difficile* was removed, the IECs washed in PBS and 2.7 ml prereduced DMEM with 10% FBS added. It was incubated for a further 3h or up to 48h. The apical and basolateral media was then removed and stored at −80°C. The IECs were washed thrice in pre-reduced PBS before lysing in 1 ml sterile water. Serial dilutions prepared from the IEC lysates were performed and plated on appropriate plates to determine the number of cell-associated bacteria.

### 4.5 Spore and total cell counts

To determine the number of spores, the lysed cells and apical supernatants were heat treated at 65°C for 20 mins as previously described (Fimlaid et al., 2015). Untreated and heat treated samples were serial diluted and plated on BHI and BHI-T agar (supplemented with 0.1% sodium taurocholate, Sigma-Aldrich, UK). No bacteria were obtained from the heat-treated samples plated on BHI (without sodium taurocholate). The CFU/ml obtained from heat treated samples plated on BHI-T plates represent heat-resistant spores, and the CFU/ml obtained from untreated samples plated on BHI plates represent the total cell counts.

### 4.6 Co-culture experiments

For the co-culture experiments with *B. dorei*, both strains were grown to stationary phase overnight in Schaedler anaerobic broth (Oxoid, UK), centrifuged with pellets resuspended in DMEM-10. OD_600_ of the suspensions were measured but not used to normalize cultures, as we found that for B. dorei the OD_600_ does not correlate well with CFU/ml. Both cultures were diluted 1:1 in DMEM-10 before loading into the VDC. The CFU/ml of these diluted cultures were determined to ensure that equal numbers of *C. difficile* to *B. dorei* (5 × 10^7^- 2.9 × 10^8^ CFU/ml) were present. Equal volumes of the diluted cultures were mixed prior to loading into the VDC. An MOI of ∼1000: 1 was used for both bacterial species. To differentiate *C. difficile* colonies from *B. dorei*, the BHIS agar was supplemented with *C. difficile* selective supplement (Oxoid, UK).

### 4.7 ELISA assays

*C. difficile* toxin production was determined using the *C. difficile* toxin A or B kit by ELISA following the manufacturer’s instructions (tgcBIOMICS, Bingen am Rhein, Germany). Briefly, the apical supernatants were centrifuged at 2500 g for 5 min and 100 µl of the supernatant in duplicates was used for the assay. Duplicates (100 µl) of the standards (toxin A and B) provided with the kit was run in the same ELISA assay as the apical supernatants from which the amount of toxin produced was calculated. IL-8 production was also determined by analysis of basolateral supernatants from VDC’s using a human IL-8 ELISA kit (R&D systems, Minneapolis, USA) following the manufacturer’s instruction. Duplicates (100 µl) of the standards (IL-8) provided with the kit was run in the same ELISA assay as the basolateral supernatants from which the amount of IL-8 produced was calculated.

### 4.8 Electrospinning protocol

Electrospinning procedure was performed as described previously (Morris et al., 2014). Briefly, scaffolds were produced by dissolving polyethylene terephthalate (PET, from food grade drinking bottles) in equal ratio of trifluoroacetic acid and dichloromethane (1:1) to create a 10% (w/v) PET solution to produce the nanofiber scaffolds. Electrospinning processing parameters included an applied voltage of 14 kV, a tip-collector distance (TCD) of 15cm, a flow rate of 0.5 ml/h and an 18 gauge spinneret for a period of 2h. Electrospinning was conducted at ambient conditions in a ducted fume hood with fibers collected on a stainless steel rotating drum (50 rpm). Once collected, the scaffolds were stored in aluminum foil at room temperature until required. Prior to cell culture, scaffolds were positioned in 12 mm Snapwell inserts (Corning, New York, USA) by replacing the commercial membrane with the electrospun nanofiber matrix. PET scaffolds were cut into 2.5 cm diameter circles and fixed with Costar Snapwell inserts using aquarium sealant glue (King British, Beaphar Company). They were sterilized by exposing to UV at the power of 80 mJ/cm^2^ for 15 minutes on each side using a UV lamp (CL-1000 Ultraviolet Crosslinker) and soaking in 10x antibiotic-antimycotic solution (final concentration: 1,000 units/ml penicillin, 1 mg/ml streptomycin and 2.5 µg/ml amphotericin B, Sigma Aldrich, UK) for 3h at 25 °C. Antibiotic-antimycotic solution was then removed and the scaffolds were washed with PBS (pH 7.4) three times. The scaffolds were air-dried in the microbiological class 2 safety cabinet overnight.

### 4.9 Scanning Electron Microscopy (SEM)

The fiber morphology of the nanofiber scaffolds was assessed using SEM. Acellular samples were sputter coated (Leica EMSCD005) with gold for 4-5 mins prior to microscopy. Scaffolds seeded with cells were fixed in 3% (v/v) glutaraldehyde overnight at 4°C. Samples were washed thrice in PBS and dehydrated through a series of industrial methylated spirits (IMS) concentrations diluted in water (25-100% v/v) for 10 mins each. Following dehydration, hexamethyldisilazane (HMDS) was added to chemically dry the samples, this process was then repeated and the samples allowed to air dry overnight before sputter coating as described above. Scaffold and cell morphology was observed at various magnifications as indicated on the scale bar of the images (JEOL JSM-6100, JEOL, UK). Images were processed using ImageJ software (W. Rasband, National Institute of Health, USA) to determine the fiber diameter and 150 measurements (50 measurements per scaffold) from 3 independently produced scaffolds were assessed.

### 4.10 Immunofluorescence assays and imaging

The Snapwell inserts, containing either the commercial membrane or PET nanofiber scaffold, were fixed in 4% PFA for 15 min, washed in PBS, permeabilized with 1% saponin (Sigma-Aldrich, UK) in 0.3% triton X-100 (Fisher Scientific, UK) and thereafter blocked against non-specific antibody binding with 3% BSA in PBS (Sigma-Aldrich, UK). For staining *C. difficile*, infected cells were incubated with anti-*C. difficile* serum for 1h, followed by Alexa Fluor 488 goat anti-rabbit secondary antibody (New England Biolabs, UK) for 1h. Alexa fluor phalloidin 647 was used to stain the actin cytoskeleton and cell nuclei were stained with ProLong Gold Antifade mountant containing 4,6-diamidino-2-phenylindole (DAPI) (New England Biolabs, UK). To determine mucus production, the Snapwell inserts were fixed, permeabilized and blocked as described above.

The Snapwell insert was incubated with mucin 2 anti-rabbit primary antibody (Santa Cruz Biotechnology, US) overnight at 4°C followed by Alexa Fluor 488 goat anti-rabbit secondary antibody (New England Biolabs, UK) for 1h. Images were taken using a confocal spinning-disk microscope (VOX UltraView, PerkinElmer, UK) with a 40X oil objective and two Hamamatsu ORCA-R2 cameras, by Volocity 6.0 (PerkinElmer, UK). Post-image analysis was performed with ImageJ software.

### 4.11 Colocalisation analysis

Coloc 2 from the ImageJ software was used to determine the colocalisation between *C. difficile*, visualized in green and actin in red, from 4 independent images at 24h and 48h p.i. Coloc 2 runs several intensity based colocalisation tests such as Manders correlation (Manders, 1993), Li intensity correlation quotient (ICQ) (Li et al., 2004) and Costes significance test (Costes et al., 2004). Intensity based analysis such as Manders’ showed the level of colocalisation between the two channels (channel 1 for actin and 2 for *C. difficile*). Manders’ output two results, M1 and M2 describe the colocalisation coefficients of the two channels. Manders’ M1 value showed that an average of 50% actin colocalised with *C.difficile* and Manders’ M2 value indicated that 100% of the *C. difficile* colocalised with actin at 24h p.i (values between 0.5 - 1). At 48h p.i, Manders M1 and M2 values showed that 100% of the actin colocalised with *C. difficile*. After applying an automatic threshold determined by the software, the levels of colocalisation was lower (Table S1). One hundred randomizations were used to determine the Costes significant P value. The Costes P value if > 95% or 0.95 was deemed to be significant. For all analyses except the Li ICQ, 1 represents perfect colocalisation with lesser values indicating the various degree of colocalisation. For the Li ICQ, the values range from maximum 0.5 to −0.5, i.e. with random (or mixed) staining ICQ= ∼0; dependent staining 0 < ICQ ≤ +0.5, and for segregated staining 0 > ICQ ≥ - 0.5 (Li et al., 2004).

### 4.12 Statistical analysis

One-way or two-way anova was used to compare two or more groups when there was one or more independent variables respectively with Tukey’s test for multiple comparison using Graphpad Prism. Student t test (two tailed) was used to determine the significance between two groups. Significance is represented as \**p* < 0.05, \*\**p* < 0.01, \*\*\**p* < 0.001 and \*\*\*\**p* < 0.0001. Except stated otherwise, all the results presented are the average of 3 independent experiments performed in duplicates or triplicates. The error bars indicate the standard deviation (SD).

## ACKNOWLEDGEMENTS

We thank Trevor Lawley for providing *C. difficile* strains and Nathalie Juge for providing the HT29-MTX cell line. We thank Arnaud Kengmo Tchoupa for reading the manuscript.

## Author contributions statement

BA, JH, JP performed experiments for this study. UD and AA produced the nanofiber scaffolds. MU, FR and SS were involved in designing experiments in the study. BA and MU wrote the main manuscript and all authors reviewed the manuscript.

## Funding

This work is funded by the Wellcome Trust Seed Award in Science to MU (grant no 108263/Z/15/Z) and a Synthetic Biology PhD funded by BBSRC & EPSRC, grant number: EP/L016494/1 to JH. This grant was awarded to the Warwick Integrative Synthetic Biology Centre (grant ref: BB/M017982/1) funded under the UK Research Councils’ Synthetic Biology for Growth programme.

## Conflict of interest statement

The authors declare no competing interests.

---

**Figure S1. Characterization of mucus production in the Caco-2/HT29-MTX monolayer.** Immunofluorescent microscopy images of Caco2/HT29-MTX monolayer (control, uninfected) stained with mucin 2 antibody showing mucus production (green) after 14 days of cell culture in the Snapwell insert. Actin is stained red and cell nuclei, blue. Scale bar = 10 µM.

**Figure S2. The vertical diffusion chamber maintains anaerobic conditions.** Overnight cultures of *C. difficile* were centrifuged and equal numbers of bacteria (3.6 × 10^7^ CFU/ml) were resuspended in DMEM-10 and incubated in the VDC or anaerobic cabinet for 3h and 24h. Serial dilutions of the apical supernatant were plated to enumerate colony counts. Data shown is the mean of 3 independent experiments and error bars indicate SD, *p* < 0.05, ns, not significant by one-way ANOVA).

**Figure S3: Change in TEER during *C. difficile* infection** Reduction in TEER measurements at different times after infection. \**p* < 0.05, \*\**p* < 0.01 and \*\*\**p* < 0.001as determined by one-way ANOVA.

**Figure S4. Spore counts from apical chamber culture supernatants from monolayer model** (A) Colony counts of spores and total cells in the apical compartment supernatants. A significant decrease in total cell count was seen at 24h and 48h p.i. but an increase in spores compared to the total bacterial numbers is observed. Data shown is the mean of 3 independent experiments and error bars indicate SD, *p* < 0.05 as determined by two-way ANOVA.

**Figure S5. Fiber diameter of the electrospun nanofibrous matrix.** Fiber diameter analysis revealed that the average fiber diameter was 457 ± 170 nm, with fibers in the range of 200-1100 nm (n = 150 measurements, 50 measurements per scaffold from 3 independently produced scaffolds).

**Figure S6. *C. difficile* spores and toxin production, and host response to infection in the multilayer model.** (A) Colony counts of spores and total cells in the host cell-associated *C. difficile* fraction of the multilayer model. Data shown is the mean of 3 independent experiments and error bars indicate SD, ns, not significant as determined by two-way ANOVA. (B) Toxin A and B levels in the multilayer model. Data shown is the mean of 3 independent experiments and error bars indicate SD, ns, not significant as determined by two-way ANOVA. Dashed line represents the sensitivity of the test at 0.5 ng/ml (C) Human IL-8 levels in the multilayer model. Data shown is the mean of 3 independent experiments and error bars indicate SD, ns, not significant as determined by the one-way ANOVA.

**Figure S7. Filaments are not produced when *C. difficile* is grown for 48h in culture in the absence of intestinal epithelial cells.** (A). *C. difficile* was grown for 48h using DMEM-10 and stained with anti-*C. difficile* for 1h, followed by Alexa Fluor 488 goat anti-rabbit secondary antibody (green). (B) The basolateral supernatant (DMEM-10) from a 24h infection in the VDC was used to grow *C. difficile* for 48h to determine if cell released factors in the supernatant can induce filament formation. Staining with anti *C. difficile* for 1h was performed, followed by Alexa Fluor 488 goat anti-rabbit secondary antibody (green). Images were taken with a Leica DMi microscope at 100x magnification. Scalebar = 10 µM

## REFERENCES

Adamu, B.O., and Lawley, T.D. (2013). Bacteriotherapy for the treatment of intestinal dysbiosis caused by *Clostridium difficile* infection. Current Opinion in Microbiology 16(5), 596–601. doi: 10.1016/j.mib.2013.06.009.

Aktories, K., Schwan, C., and Jank, T. (2017). *Clostridium difficile* Toxin Biology. Annual Review of Microbiology 71(1), 281–307. doi: 10.1146/annurev-micro-090816-093458.

Barketi-Klai, A., Hoys, S., Lambert-Bordes, S., Collignon, A., and Kansau, I. (2011). Role of fibronectin-binding protein A in *Clostridium difficile* intestinal colonization. Journal of Medical Microbiology 60(8), 1155–1161. doi: doi:10.1099/jmm.0.029553-0.

Barrila, J., Yang, J., Crabbé, A., Sarker, S.F., Liu, Y., Ott, C.M., et al. (2017). Three-dimensional organotypic co-culture model of intestinal epithelial cells and macrophages to study *Salmonella enterica* colonization patterns. npj Microgravity 3(1), 10. doi: 10.1038/s41526-017-0011-2.

Batah, J., Kobeissy, H., Bui Pham, P.T., Deneve-Larrazet, C., Kuehne, S., Collignon, A., et al. (2017). *Clostridium difficile* flagella induce a pro-inflammatory response in intestinal epithelium of mice in cooperation with toxins. Sci Rep 7(1), 3256. doi: 10.1038/s41598-017-03621-z.

Bobo, L.D., El Feghaly, R.E., Chen, Y.S., Dubberke, E.R., Han, Z., Baker, A.H., et al. (2013). MAPK-activated protein kinase 2 contributes to Clostridium difficile-associated inflammation. Infect Immun 81(3), 713–722. doi: 10.1128/IAI.00186-12.

Calabi, E., Calabi, F., Phillips, A.D., and Fairweather, N.F. (2002). Binding of *Clostridium difficile* Surface Layer Proteins to Gastrointestinal Tissues. Infection and Immunity 70(10), 5770–5778. doi: 10.1128/iai.70.10.5770-5778.2002.

Carter, G.P., Chakravorty, A., Pham Nguyen, T.A., Mileto, S., Schreiber, F., Li, L., et al. (2015). Defining the Roles of TcdA and TcdB in Localized Gastrointestinal Disease, Systemic Organ Damage, and the Host Response during Clostridium difficile Infections. MBio 6(3), e00551. doi: 10.1128/mBio.00551-15.

Cerquetti, M., Serafino, A., Sebastianelli, A., and Mastrantonio, P. (2002). Binding of *Clostridium difficile* to Caco-2 epithelial cell line and to extracellular matrix proteins. FEMS Immunology & Medical Microbiology 32(3), 211–218. doi: 10.1111/j.1574-695X.2002.tb00556.x.

Chen, S., Sun, C., Wang, H., and Wang, J. (2015). The Role of Rho GTPases in Toxicity of Clostridium difficile Toxins. Toxins (Basel) 7(12), 5254–5267. doi: 10.3390/toxins7124874.

Chen, X., Katchar, K., Goldsmith, J.D., Nanthakumar, N., Cheknis, A., Gerding, D.N., et al. (2008). A mouse model of Clostridium difficile-associated disease. Gastroenterology 135(6), 1984–1992. doi: 10.1053/j.gastro.2008.09.002.

Collins, D.A., Selvey, L.A., Celenza, A., and Riley, T.V. (2017). Community-associated *Clostridium difficile* infection in emergency department patients in Western Australia. Anaerobe 48(Supplement C), 121–125. doi: https://doi.org/10.1016/j.anaerobe.2017.08.008.

Costes, S.V., Daelemans, D., Cho, E.H., Dobbin, Z., Pavlakis, G., and Lockett, S. (2004). Automatic and quantitative measurement of protein-protein colocalization in live cells. Biophys J 86(6), 3993–4003. doi: 10.1529/biophysj.103.038422.

Darkoh, C., Odo, C., and DuPont, H.L. (2016). Accessory Gene Regulator-1 Locus Is Essential for Virulence and Pathogenesis of Clostridium difficile. MBio 7(4). doi: 10.1128/mBio.01237-16.

Davies, K.A., Longshaw, C.M., Davis, G.L., Bouza, E., Barbut, F., Barna, Z., et al. (2014). Underdiagnosis of *Clostridium difficile* across Europe: the European, multicentre, prospective, biannual, point-prevalence study of *Clostridium difficile* infection in hospitalised patients with diarrhoea (EUCLID). The Lancet Infectious Diseases 14(12), 1208–1219. doi: 10.1016/S1473-3099(14)70991-0.

Deakin, L.J., Clare, S., Fagan, R.P., Dawson, L.F., Pickard, D.J., West, M.R., et al. (2012a). The *Clostridium difficile* spo0A Gene Is a Persistence and Transmission Factor. Infection and Immunity 80(8), 2704–2711. doi: 10.1128/iai.00147-12.

Deakin, L.J., Clare, S., Fagan, R.P., Dawson, L.F., Pickard, D.J., West, M.R., et al. (2012b). Clostridium difficile spo0A gene is a persistence and transmission factor. Infect Immun. doi: 10.1128/IAI.00147-12.

DeCicco RePass, M.A., Chen, Y., Lin, Y., Zhou, W., Kaplan, D.L., and Ward, H.D. (2017). Novel Bioengineered Three-Dimensional Human Intestinal Model for Long-Term Infection of *Cryptosporidium parvum*. Infection and Immunity 85(3), e00731–00716. doi: 10.1128/IAI.00731-16.

Drummond, C.G., Nickerson, C.A., and Coyne, C.B. (2016). A Three-Dimensional Cell Culture Model To Study Enterovirus Infection of Polarized Intestinal Epithelial Cells. mSphere 1(1). doi: 10.1128/mSphere.00030-15.

England, P.H. (2017). Annual Epidemiological Commentary: Mandatory MRSA, MSSA and E. coli bacteraemia and C. difficile infection data, 2016/17.

Fimlaid, K.A., Jensen, O., Donnelly, M.L., Siegrist, M.S., and Shen, A. (2015). Regulation of Clostridium difficile Spore Formation by the SpoIIQ and SpoIIIA Proteins. PLoS Genet 11(10), e1005562. doi: 10.1371/journal.pgen.1005562.

Fuentes, S., van Nood, E., Tims, S., Heikamp-de Jong, I., ter Braak, C.J.F., Keller, J.J., et al. (2014). Reset of a critically disturbed microbial ecosystem: faecal transplant in recurrent *Clostridium difficile* infection. The ISME Journal 8(8), 1621–1633. doi: 10.1038/ismej.2014.13.

Gravel, D., Miller, M., Simor, A., Taylor, G., Gardam, M., McGeer, A., et al. (2009). Health Care-Associated Clostridium difficile Infection in Adults Admitted to Acute Care Hospitals in Canada: A Canadian Nosocomial Infection Surveillance Program Study. Clin Infect Dis 48. doi: 10.1086/596703.

Hennequin, C., Porcheray, F., Waligora-Dupriet, A.-J., Collignon, A., Barc, M.-C., Bourlioux, P., et al. (2001). GroEL (Hsp60) of *Clostridium difficile* is involved in cell adherence. Microbiology 147(1), 87–96. doi: 10.1099/00221287-147-1-87.

Hong, H.A., Ferreira, W.T., Hosseini, S., Anwar, S., Hitri, K., Wilkinson, A.J., et al. (2017). The Spore Coat Protein CotE Facilitates Host Colonization by Clostridium difficile. J Infect Dis 216(11), 1452–1459. doi: 10.1093/infdis/jix488.

Huttenhower, C., Gevers, D., Knight, R., Abubucker, S., Badger, J.H., Chinwalla, A.T., et al. (2012). Structure, function and diversity of the healthy human microbiome. Nature 486(7402), 207.

Jafari, N.V., Kuehne, S.A., Minton, N.P., Allan, E., and Bajaj-Elliott, M. (2016). *Clostridium difficile*-mediated effects on human intestinal epithelia: Modelling host-pathogen interactions in a vertical diffusion chamber. Anaerobe 37(Supplement C), 96–102. doi: https://doi.org/10.1016/j.anaerobe.2015.12.007.

Janvilisri, T., Scaria, J., and Chang, Y.-F. (2010). Transcriptional profiling of *Clostridium difficile* and Caco-2 cells during infection. The Journal of infectious diseases 202(2), 282–290. doi: 10.1086/653484.

Justice, S.S., Hunstad, D.A., Cegelski, L., and Hultgren, S.J. (2008). Morphological plasticity as a bacterial survival strategy. Nat Rev Micro 6(2), 162–168.

Kalita, A., Hu, J., and Torres, A.G. (2014). Recent advances in adherence and invasion of pathogenic Escherichia coli. Curr Opin Infect Dis 27(5), 459–464. doi:10.1097/QCO.0000000000000092.

Kasendra, M., Barrile, R., Leuzzi, R., and Soriani, M. (2014). *Clostridium difficile* Toxins Facilitate Bacterial Colonization by Modulating the Fence and Gate Function of Colonic Epithelium. Journal of Infectious Diseases 209(7), 1095–1104. doi: 10.1093/infdis/jit617.

Kook, Y.-M., Jeong, Y., Lee, K., and Koh, W.-G. (2017). Design of biomimetic cellular scaffolds for co-culture system and their application. Journal of Tissue Engineering 8, 2041731417724640. doi: 10.1177/2041731417724640.

Kovacs-Simon, A., Leuzzi, R., Kasendra, M., Minton, N., Titball, R.W., and Michell, S.L. (2014). Lipoprotein CD0873 Is a Novel Adhesin of *Clostridium difficile*. The Journal of Infectious Diseases 210(2), 274–284. doi: 10.1093/infdis/jiu070.

Kuehne, S.A., Cartman, S.T., Heap, J.T., Kelly, M.L., Cockayne, A., and Minton, N.P. (2010). The role of toxin A and toxin B in *Clostridium difficile* infection. Nature 467(7316), 711–713. doi: 10.1038/nature09397.

Kuehne, S.A., Collery, M.M., Kelly, M.L., Cartman, S.T., Cockayne, A., and Minton, N.P. (2013). Importance of toxin A, toxin B, and CDT in virulence of an epidemic *Clostridium difficile* strain. The Journal of infectious diseases 209(1), 83–86.

Kuehne, S.A., Collery, M.M., Kelly, M.L., Cartman, S.T., Cockayne, A., and Minton, N.P. (2014). Importance of toxin A, toxin B, and CDT in virulence of an epidemic Clostridium difficile strain. J Infect Dis 209(1), 83–86. doi: 10.1093/infdis/jit426.

Lawley, T.D., Clare, S., Deakin, L.J., Goulding, D., Yen, J.L., Raisen, C., et al. (2010). Use of Purified *Clostridium difficile* Spores To Facilitate Evaluation of Health Care Disinfection Regimens. Applied and Environmental Microbiology 76(20), 6895–6900. doi: 10.1128/aem.00718-10.

Leslie, J.L., Huang, S., Opp, J.S., Nagy, M.S., Kobayashi, M., Young, V.B., et al. (2015). Persistence and Toxin Production by *Clostridium difficile* within Human Intestinal Organoids Result in Disruption of Epithelial Paracellular Barrier Function. Infection and Immunity 83(1), 138–145. doi: 10.1128/iai.02561-14.

Lessa, F.C., Mu, Y., Bamberg, W.M., Beldavs, Z.G., Dumyati, G.K., Dunn, J.R., et al. (2015). Burden of *Clostridium difficile* infection in the United States. N Engl J Med 372. doi: 10.1056/NEJMoa1408913.

Li, Q., Lau, A., Morris, T.J., Guo, L., Fordyce, C.B., and Stanley, E.F. (2004). A syntaxin 1, Galpha(o), and N-type calcium channel complex at a presynaptic nerve terminal: analysis by quantitative immunocolocalization. J Neurosci 24(16), 4070–4081. doi: 10.1523/JNEUROSCI.0346-04.2004.

Lloyd-Price, J., Abu-Ali, G., and Huttenhower, C. (2016). The healthy human microbiome. Genome Medicine 8.

Lyras, D., O’Connor, J.R., Howarth, P.M., Sambol, S.P., Carter, G.P., Phumoonna, T., et al. (2009). Toxin B is essential for virulence of Clostridium difficile. Nature 458(7242), 1176–1179. doi: 10.1038/nature07822.

Manders, E.M., Verbeek, F. J. and Aten J.A (1993). Measurement of co-localization of objects in dual-colour confocal images. Journal of Microcopy 169(3), 375–382.

McKee, R.W., Aleksanyan, N., Garrett, E.M., and Tamayo, R. (2018). Type IV Pili Promote Clostridium difficile Adherence and Persistence in a Mouse Model of Infection. Infect Immun 86(5). doi: 10.1128/IAI.00943-17.

Merrigan, M.M., Venugopal, A., Roxas, J.L., Anwar, F., Mallozzi, M.J., Roxas, B.A.P., et al. (2013). Surface-Layer Protein A (SlpA) Is a Major Contributor to Host-Cell Adherence of *Clostridium difficile*. PLOS ONE 8(11), e78404. doi: 10.1371/journal.pone.0078404.

Mills, D.C., Gundogdu, O., Elmi, A., Bajaj-Elliott, M., Taylor, P.W., Wren, B.W., et al. (2012). Increase in *Campylobacter jejuni* Invasion of Intestinal Epithelial Cells under Low-Oxygen Coculture Conditions That Reflect the In Vivo Environment. Infection and Immunity 80(5), 1690–1698. doi: 10.1128/iai.06176-11.

Mora-Uribe, P., Miranda-Cárdenas, C., Castro-Córdova, P., Gil, F., Calderón, I., Fuentes, J.A., et al. (2016). Characterization of the Adherence of *Clostridium difficile* Spores: The Integrity of the Outermost Layer Affects Adherence Properties of Spores of the Epidemic Strain R20291 to Components of the Intestinal Mucosa. Frontiers in Cellular and Infection Microbiology 6, 99. doi: 10.3389/fcimb.2016.00099.

Morris, G.E., Bridge, J.C., Brace, L.A., Knox, A.J., Aylott, J.W., Brightling, C.E., et al. (2014). A novel electrospun biphasic scaffold provides optimal three-dimensional topography for in vitro co-culture of airway epithelial and fibroblast cells. Biofabrication 6(3), 035014. doi: 10.1088/1758-5082/6/3/035014.

Paredes-Sabja, D., and Sarker, M.R. (2012). Adherence of *Clostridium difficile* spores to Caco-2 cells in culture. Journal of Medical Microbiology 61(9), 1208–1218. doi: doi:10.1099/jmm.0.043687-0.

Pumbwe, L., Skilbeck, C.A., Nakano, V., Avila-Campos, M.J., Piazza, R.M.F., and Wexler, H.M. (2007). Bile salts enhance bacterial co-aggregation, bacterial-intestinal epithelial cell adhesion, biofilm formation and antimicrobial resistance of *Bacteroides fragilis*. Microbial Pathogenesis 43(2), 78–87. doi: https://doi.org/10.1016/j.micpath.2007.04.002.

Qin, J., Li, R., Raes, J., Arumugam, M., Burgdorf, K.S., and Manichanh, C. (2010). A human gut microbial gene catalogue established by metagenomic sequencing. Nature 464. doi: 10.1038/nature08821.

Rao, K., Erb-Downward, J.R., Walk, S.T., Micic, D., Falkowski, N., Santhosh, K., et al. (2014). The systemic inflammatory response to *Clostridium difficile* infection. PLoS One 9(3), e92578.

Ravi, M., Paramesh, V., Kaviya, S.R., Anuradha, E., and Solomon, F.D.P. (2015). 3D Cell Culture Systems: Advantages and Applications. Journal of Cellular Physiology 230(1), 16–26. doi: 10.1002/jcp.24683.

Ribet, D., and Cossart, P. (2015). How bacterial pathogens colonize their hosts and invade deeper tissues. Microbes Infect 17(3), 173–183. doi: 10.1016/j.micinf.2015.01.004.

Schüller, S., and Phillips, A.D. (2010). Microaerobic conditions enhance type III secretion and adherence of enterohaemorrhagic *Escherichia coli* to polarized human intestinal epithelial cells. Environmental Microbiology 12(9), 2426–2435. doi: 10.1111/j.1462-2920.2010.02216.x.

Shaban, L., Chen, Y., Fasciano, A.C., Lin, Y., Kaplan, D.L., Kumamoto, C.A., et al. (2018). A 3D intestinal tissue model supports Clostridioides difficile germination, colonization, toxin production and epithelial damage. Anaerobe 50, 85–92. doi: 10.1016/j.anaerobe.2018.02.006.

Smith, R.S., Smith, T.J., Blieden, T.M., and Phipps, R.P. (1997). Fibroblasts as sentinel cells. Synthesis of chemokines and regulation of inflammation. The American Journal of Pathology 151(2), 317–322.

Tasteyre, A., Barc, M.-C., Collignon, A., Boureau, H., and Karjalainen, T. (2001). Role of FliC and FliD Flagellar Proteins of *Clostridium difficile* in Adherence and Gut Colonization. Infection and Immunity 69(12), 7937–7940. doi: 10.1128/iai.69.12.7937-7940.2001.

Vohra, P., and Poxton, I.R. (2011). Efficacy of decontaminants and disinfectants against *Clostridium difficile*. Journal of Medical Microbiology 60(8), 1218–1224. doi: doi:10.1099/jmm.0.030288-0.

Waligora, A.-J., Hennequin, C., Mullany, P., Bourlioux, P., Collignon, A., and Karjalainen, T. (2001). Characterization of a Cell Surface Protein of *Clostridium difficile* with Adhesive Properties. Infection and Immunity 69(4), 2144–2153. doi: 10.1128/iai.69.4.2144-2153.2001.

Yoon, S., Yu, J., McDowell, A., Kim, S.H., You, H.J., and Ko, G. (2017). Bile salt hydrolase-mediated inhibitory effect of *Bacteroides ovatus* on growth of *Clostridium difficile*. Journal of Microbiology 55(11), 892–899. doi: 10.1007/s12275-017-7340-4.

