## Supplementary figures and images for "Probing *Clostridium difficile* infection in innovative human gut cellular models"

### Fig S1

Fig S1

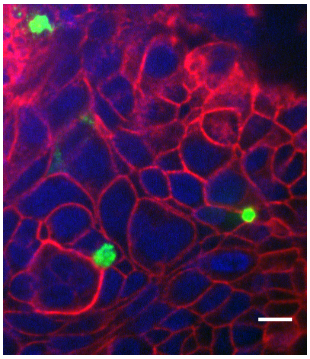

### Fig S2

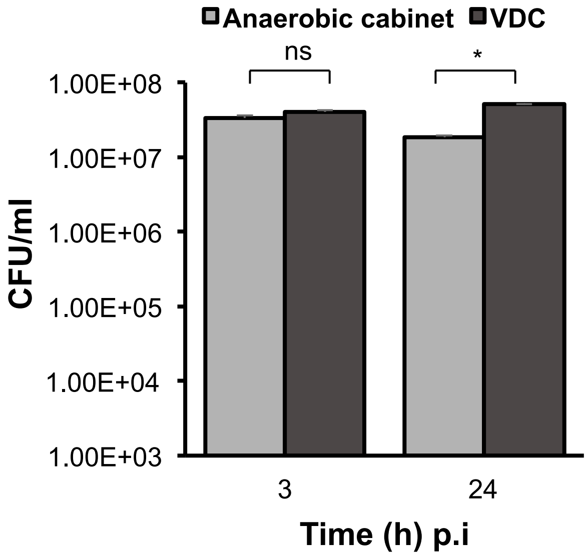

### Fig S3

#### Figure S3

A

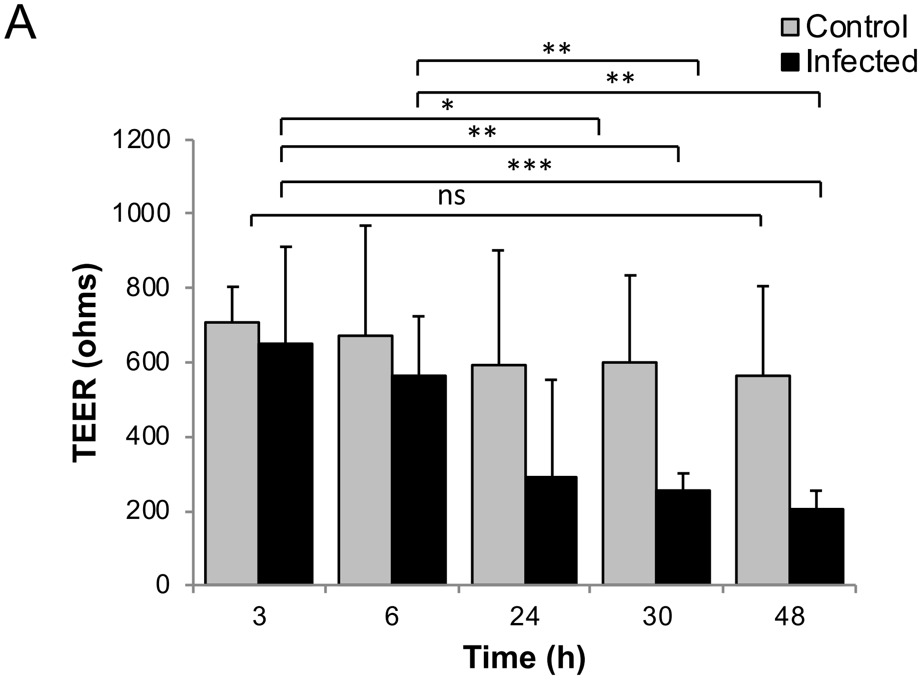


B

WT 24h p.i.

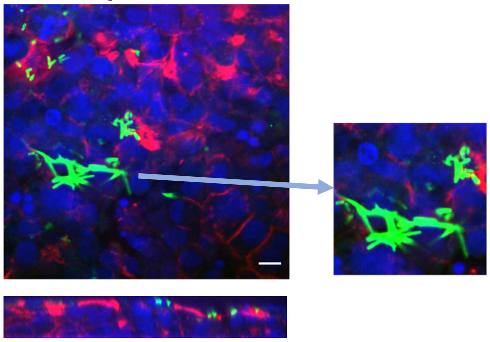

### Fig S4

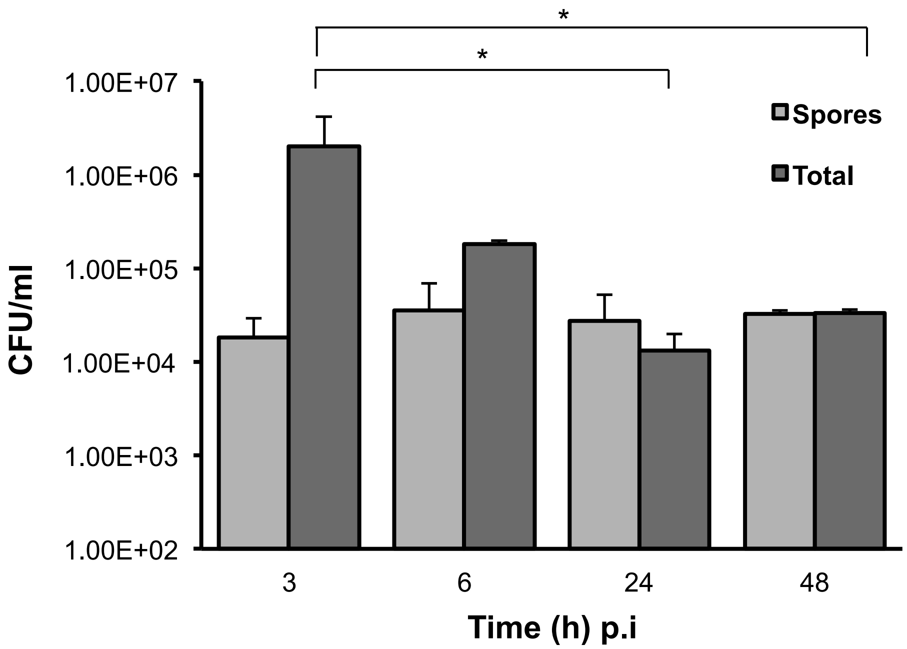

### Fig S5

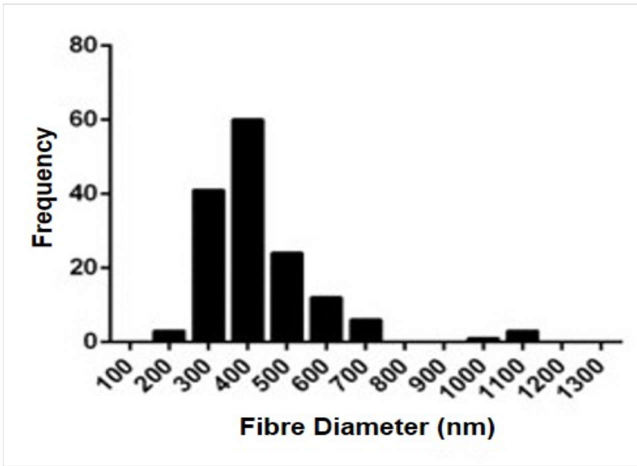

### Fig S6

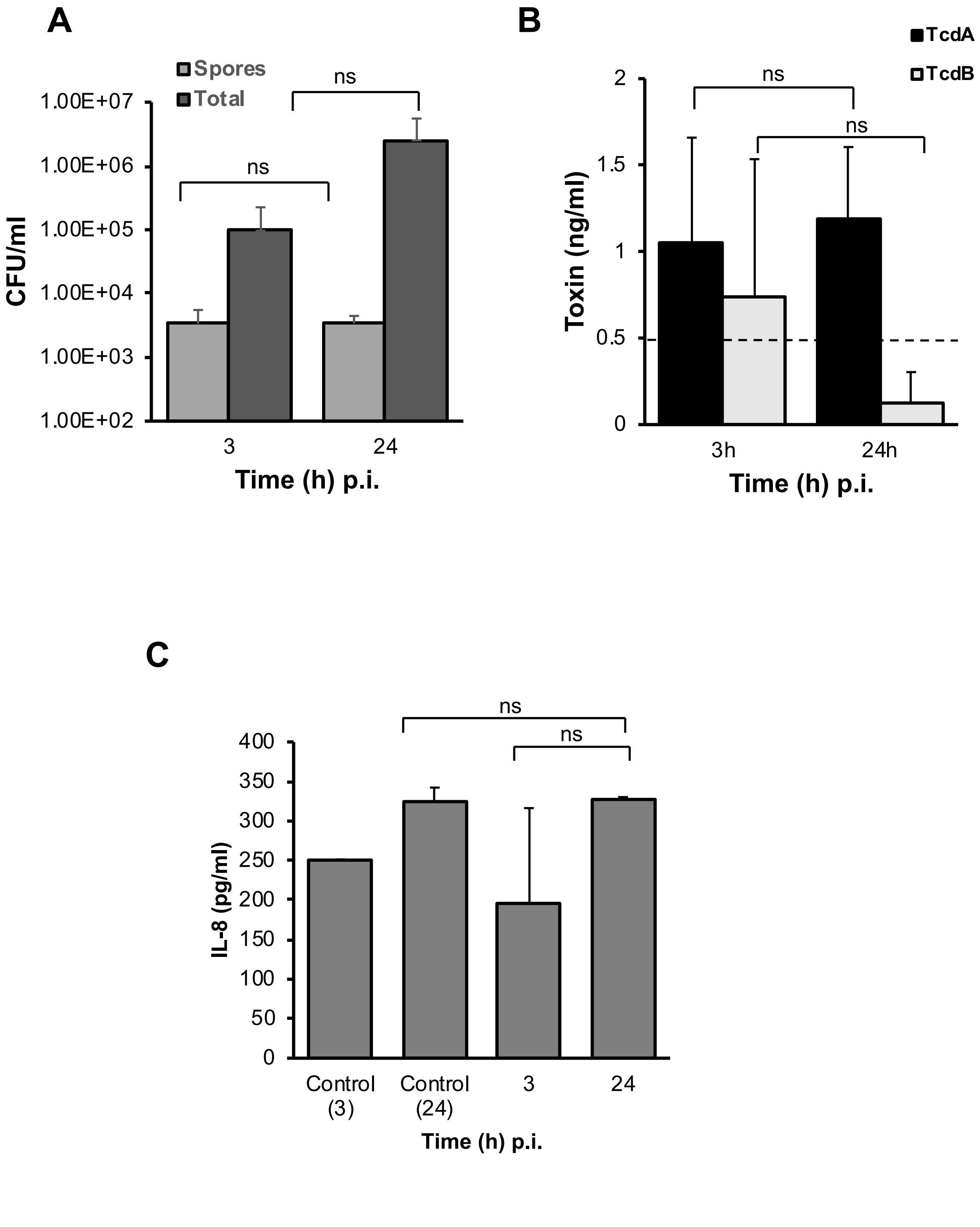

### Fig S7

Figure S7

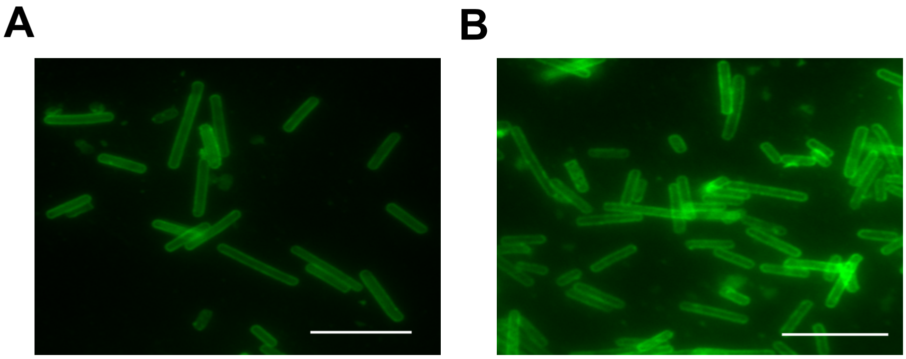
